# Mer3 helicase protects early crossover intermediates from STR complex disassembly during meiosis

**DOI:** 10.1101/2022.06.27.497840

**Authors:** Veronika Altmannova, Magdalena Firlej, Franziska Müller, Petra Janning, Rahel Rauleder, Dorota Rousova, Andreas Schäffler, John R. Weir

## Abstract

During meiosis I it is necessary that homologous chromosomes are linked to one another so that they can be faithfully separated. *S. cerevisiae* Mer3 (HFM1 in mammals) is a SF2 helicase and member of the ZMM group of proteins, that facilitates the formation of class I crossovers during meiosis. Here we describe the structural organisation of Mer3 and, using AlphaFold modelling and XL-MS, we further characterise the previously described interaction with Mlh1-Mlh2. We find that Mer3 also forms a previously undescribed complex with the recombination regulating factors Top3 and Rmi1 and that this interaction is competitive with Sgs1^BLM^ helicase in a phospho-dependent manner. Using *in vitro* reconstituted D-loop assays we show that Mer3 inhibits the anti-recombination activity of Sgs1/Top3/Rmi1 (STR) complex. Thus we provide a mechanism whereby Mer3 downregulates the anti-crossover activity of the STR complex, hence promoting the formation of crossovers during meiosis I.

## Introduction

Most sexually reproducing organisms utilise meiotic recombination to both link homologous chromosomes during meiosis I, and to generate genetic diversity among their gametes and subsequent progeny. Recombination is initiated by the controlled generation of double-stranded DNA breaks at the onset of prophase I (reviewed in Lam and Keeney, 2014 and Yadav and Claeys Bouuaert, 2021). While the initial processing of these breaks is analogous to DNA repair in the soma, two important modulations occur in the germline to generate the required crossovers (COs) between homologous chromosomes. Firstly, repair must take place from the homologous chromosome rather than the sister (inter homolog bias, reviewed in Humphryes and Hochwagen, 2014), secondly the repair intermediates must be protected from disassembly by anti-CO factors, that prevent the formation of inter-homolog COs and a resulting loss of heterozygosity (LoH) in the soma (LaRocque *et al*., 2011).

One such “anti-crossover” factor is the *S. cerevisiae* helicase Sgs1, which is functionally orthologous to the Bloom-syndrome helicase BLM (Bernstein *et al*., 2010). Sgs1 performs its activities in a complex with the type IA topoisomerase Top3, and the OB-fold accessory factor Rmi1 (Bennett *et al*., 2000; Bernstein *et al*., 2010; Johnson *et al*., 2000; Mullen *et al*., 2005). STR complex combines helicase and decatenase activities to displace strand invasion intermediates (Bachrati *et al*., 2006; van Brabant *et al*., 2000), and dissolve double-Holliday junctions (dHJs) (Bizard and Hickson, 2014), and thus contributes to genome stability in mitotically dividing cells. During meiosis Sgs1 and STR activity is, somewhat counter intuitively, required for normal crossover formation (Amin *et al*., 2010; De Muyt *et al*., 2012; Oh *et al*., 2007; Tang *et al*., 2015). However, Sgs1 is not always active during meiosis, and its activity is instead promoted through CDK phosphorylation, leading to a temporal separation of pro- and anti-crossover activities (Grigaitis *et al*., 2020).

In budding yeast, a group of proteins was identified that, in general, promoted CO formation, and was collectively termed “ZMM” (Brner *et al*., 2004) The *S. cerevisiae* group of ZMM proteins consists of Zip1, Zip2, Zip3, Zip4, Spo16, the Mer3 helicase, and the Msh4-Msh5 heterodimer (Lynn *et al*., 2007; Shinohara *et al*., 2008). Some ZMMs are involved in the formation and stabilisation of single-end invasion (SEI) intermediates (Brner *et al*., 2004; Hunter and Kleckner, 2001); ZMM mutants show a decrease in the formation of SEI and dHJ intermediates. In absence of ZMMs spore viability is decreased as well as the number of COs (Brner *et al*., 2004; Jessop *et al*., 2006). ZMM proteins were also presumed to downregulate the activity of Sgs1 helicase (Jessop *et al*., 2006).

Mer3 helicase is well conserved, with functional orthologs being found in other fungi (Storlazzi *et al*., 2010), plants (Chen *et al*., 2005; Mercier *et al*., 2005; Wang *et al*., 2009) and mammals (Guiraldelli *et al*., 2013), where it is also required for human fertility (Wang *et al*., 2014). In vitro studies on Mer3 showed that it is an active ATPase with strand separation activity working in 3’ to 5’ direction (Nakagawa *et al*., 2001) and that it might preferentially recognize Holliday junctions, however, it also recognizes other DNA structures (Duroc *et al*., 2017; Nakagawa and Kolodner, 2002). Further in vitro works demonstrated that Mer3 promoted a heteroduplex extension by Rad51, that is, it enlarged and stabilised D-loops (Mazina *et al*., 2004). It was shown that Mer3, together with other ZMM proteins, synergize to protect nascent CO-designated recombination intermediates from disassembly by Sgs1 (Jessop *et al*., 2006). While the mechanisms for chromosomal recruitment of Mer3 are unclear, it does appear that Mer3 is recruited early in the DNA repair pathway (Pyatnitskaya *et al*., 2019; Storlazzi *et al*., 2010). In vivo Mer3 ATPase deficient mutants (mer3G166D and mer3K167A) show mild spore viability defects whereas in mer3Δ strain spore viability is strongly compromised (Nakagawa and Kolodner, 2002). This observation hints at the possibility that Mer3 may contribute to promoting crossover formation through protein-protein interactions. To date Mer3 has been reported to interact with only a few proteins involved in meiotic recombination such as the helicase Pif1, replication factor Rfc1, (Vernekar *et al*., 2021), and a MutLβ complex (Mlh1/Mlh2) (Duroc *et al*., 2017). The interaction between Mer3 and MutLβ was shown to occur via the Ig-like domain of Mer3 (Duroc *et al*., 2017). Impairing the ability of Mer3 to bind to Mlh2 leads to an increase in the length of gene conversion tracts, both in COs and in NCOs (Duroc *et al*., 2017).

Here we used biochemical and structural approaches to describe the activity, oligomeric status and structural organisation of Mer3. We also further characterised the details of the Mer3 interaction with the Mlh1/Mlh2 complex. The search for novel Mer3 interactors has led us to discover that Mer3 forms a complex with Top3 and Rmi1 and that this interaction is compatible with Mlh1/Mlh2 binding. Interestingly, Mer3 is able to disrupt Top3/Rmi1 binding to Sgs1BLM helicase in a phospho-dependent manner. We further show that Mer3 inhibits D-loop disassembly mediated by the Sgs1-Top3/Rmi1 complex thus revealing a novel mechanism for the protection of DNA repair intermediates and thus promotion of crossover formation during meiosis I.

## Results

### Hybrid structural and biophysical analysis of Mer3

We set out by purifying full-length *S. cerevisiae* Mer3 from baculovirus-infected insect cells using a COOH-terminal 2xStrep-II tag. Using a 3-step purification (see materials and methods) we were able to produce a Mer3 that was homogenous and devoid of nucleic acid contamination (Figure 1B). Using mass photometry, we determined our protein preparation to be a homogenous sample of monomers of Mer3 in the solution at a concentration of 30 nM (Figure 1C). We tested the DNA binding of recombinant Mer3 and found that it bound both single-stranded DNA (ssDNA) and synthetic “D-loop” substrates with high affinity (Supplementary Figure 1A). Tight binding to D-loops is consistent with previous studies (Duroc *et al*., 2017; Mazina *et al*., 2004), but high affinity for ssDNA was not reported to date. We also confirmed that our Mer3 preparation was catalytically active in a strand-separation assay (Supplementary Figure 1B).

**Figure 1.**
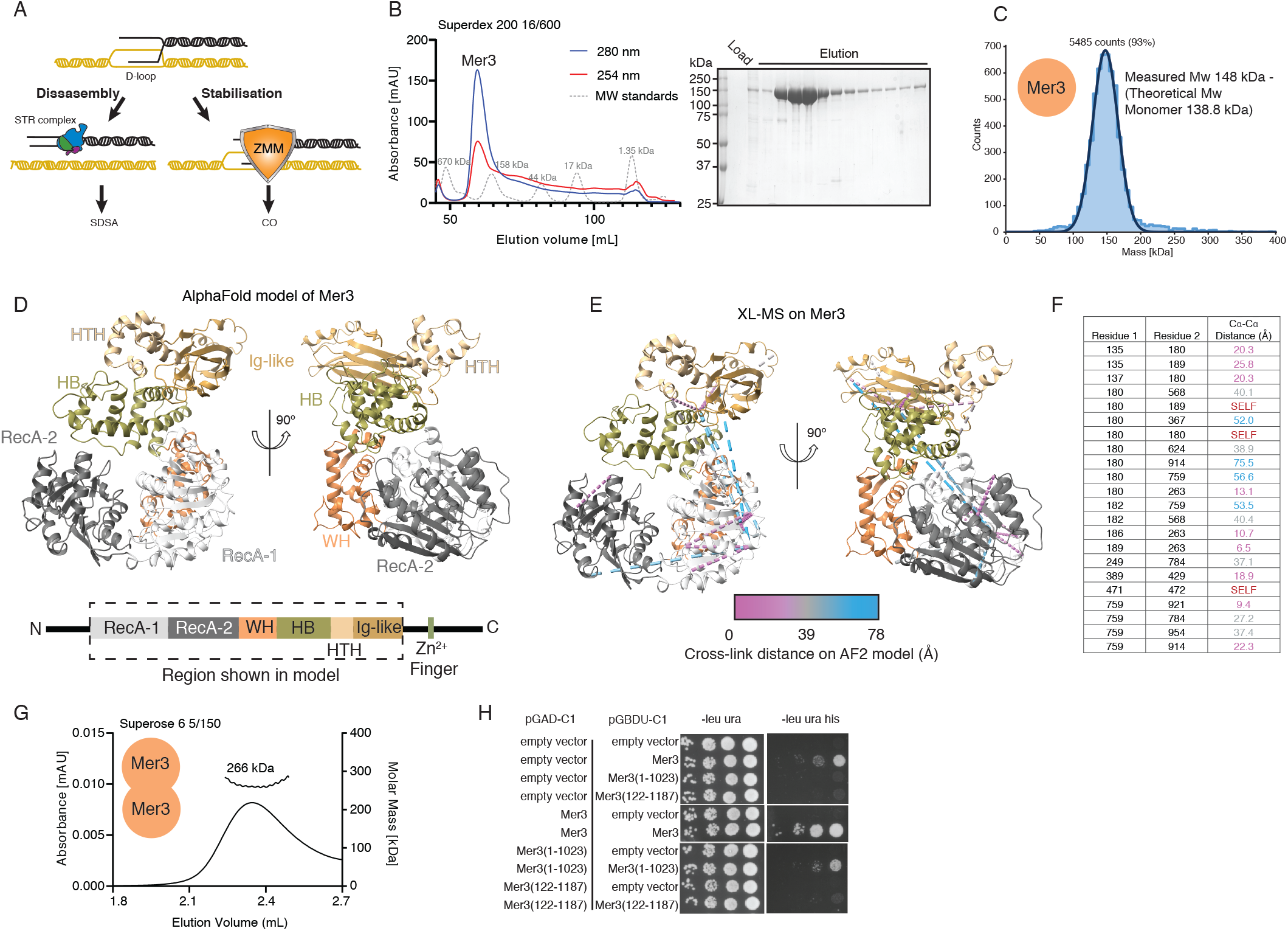
Biophysical and Structural Analysis of Mer3 hellicase. A) Cartoon representation of two possible outcomes for D-loop DNA repair intermediates during meiosis in budding yeast. Either the D-loop is disassembled by the Sgs1 helicase (in the Sgs1/Top3/Rmi1 - STR - complex) or the D-loop is captured and stabilised by members of the ZMM group of proteins B) Purification of the meiosis-specific helicase Mer3. Upper panel - size exclusion chromatography (Superdex 200 16/600) profile of recombinant Mer3; red trace = absorbance at 254 nm, blue trace absorbance at 280 nm, grey trace = SEC standards. Lower panel - coomassie stained SDS-PAGE gel of the SEC column elution. C) Molecular mass determination of Mer3 using Mass Photometry (30 nM protein concentration), showing that we only detect Mer3 monomers as this concentration. D) AlphaFold2 model of S. cerevisiae Mer3 from the publicly available database (https://www.alphafold.ebi.ac.uk/). Domains are assigned based on the Brr2 helicase helicase structure (Pena et al., 2009). E) Cross-links (DSBU cross-linker) modelled onto the Mer3 AF2 structure, and coloured according to distance using XMAS (Lagerwaard et al., 2022), with purple cross-links generally consistent with the model, and blue cross-links in violation. F) List of Mer3 cross-links indicated the residues cross-links and the range of distances (cross-links of <30an be considered compatible with the model). The self cross-links are highlighted that can only be compatible with a higher order complex. G) SEC-MALS on Mer3 (10 M injection concentration) showing that Mer3 can form homodimers at higher concentrations. H) Y2H analysis of the Mer3 self-association regions.

To probe the structure of Mer3, we made use of the AlphaFold2 (AF2) (Jumper *et al*., 2021) predicted model that is publicly available (AlphaFold EBI ID P51979). Based on the pLDDT score and the predicted error alignment (PAE) plots the AF2 model of Mer3 is of an overall high quality (Supplementary Figure 1C and D). The predicted model reveals an architecture consisting of (from N- to C-terminus) RecA-like 1, RecA-like 2, winged helix (WH), helical bundle (HB), helix-loop-helix (HLH) and Ig-like domains (Figure 1D). A DALI (Holm, 2020) search reveals the greatest similarity to the spliceosomal helicase Brr2 (Pena *et al*., 2009), in particular the presence of the HB, HLH and Ig-like domains in what has been previously described as a Sec63 like-region (Pena *et al*., 2009; Ponting, 2000).

We validated the AF2 model prediction using chemical cross-linking coupled to mass-spectrometry (XL-MS). We used DSBU (Mller *et al*., 2010; Pan *et al*., 2018) as a bifunctional chemical cross-linker to produce an inter-residue distance map of Mer3 (Supplementary Figure 1E). We modelled the cross-links onto the AF2 model of Mer3, and determined which cross-links were consistent with the inter-residue distances in the AF2 model, and which were violated. Surprisingly, despite the high quality of the AF2 prediction, a proportion of Mer3 observed cross-link distances were not compatible with the model (distances of >30Å, Figure 1E and 1F). Closer analysis of the cross-linking data revealed a number of “self” cross-linked peptides that could only be compatible with a higher order stoichiometry (Figure 1F). This is superficially surprising, since we had already demonstrated that Mer3 is a monomer (Figure 1B), however at a 100-fold lower concentration than we used for XL-MS (30 nM in the mass photometer vs. 3 μM in the reaction with DSBU). Since a sample concentration of 3 μM is not currently compatible with the mass-photometry method, we instead used multi-angle light scattering coupled to size-exclusion chromatography (SEC-MALS) to measure the molecular mass of Mer3 at higher (10 μM at injection) concentrations. SEC-MALS revealed a single species with a molecular mass of 266 kDa, consistent with a dimer of Mer3 (theoretical molecular mass 277.6 kDa) (Figure 1G). As such we conclude that Mer3 forms dimers at higher concentrations with a KD in the high nM to low μM range.

To map the possible oligomerization region we analysed Mer3 truncations. Constructs of Mer3 that disrupted the structural core of Mer3 were unstable in recombinant preparations. Instead we made use of constructs lacking the N- or C-terminal unstructured regions: Mer3ΔN (122-1187), Mer3ΔC (1-1023), based on the AF2 prediction. We carried out initial tests using yeast-two-hybrid (Figure 1H) which suggested that both the N- and C-termini contributed to self association. As such we conclude that Mer3 forms dimers at higher concentrations, and that this dimerisation requires both the N- and C-terminal unstructured regions, possibly indicating at least in part a trans N- to C-terminal interaction.

### Biophysical and structural analysis of Mer3 interaction with Mlh1 and Mlh2

It was previously shown that Mer3 binds directly to the Mlh1/Mlh2 complex and that this interaction plays a role in regulating the size of gene conversions during NCOs (Duroc *et al*., 2017). To investigate the structural organisation of this complex we purified a complex of MBP-Mlh1 and MBP-Mlh2 from insect cells, again making use of a 3-step purification to ensure that it was free of nucleic acids (Figure 2A, Supplementary figure 2A). Both proteins were 6xHis-MBP tagged because purification of these proteins without the solubilization tag resulted in a much lower yield and in our experiments, we also left the MBP-tag on both proteins for the downstream experiments due to continued problems with solubility. Using mass photometry we determined the molecular mass of the purified Mlh1/Mlh2 complex to be 289 kDa; consistent with a heterodimer (theoretical mass of a heterodimer = 250.8 kDa) (Figure 2B).

**Figure 2.**
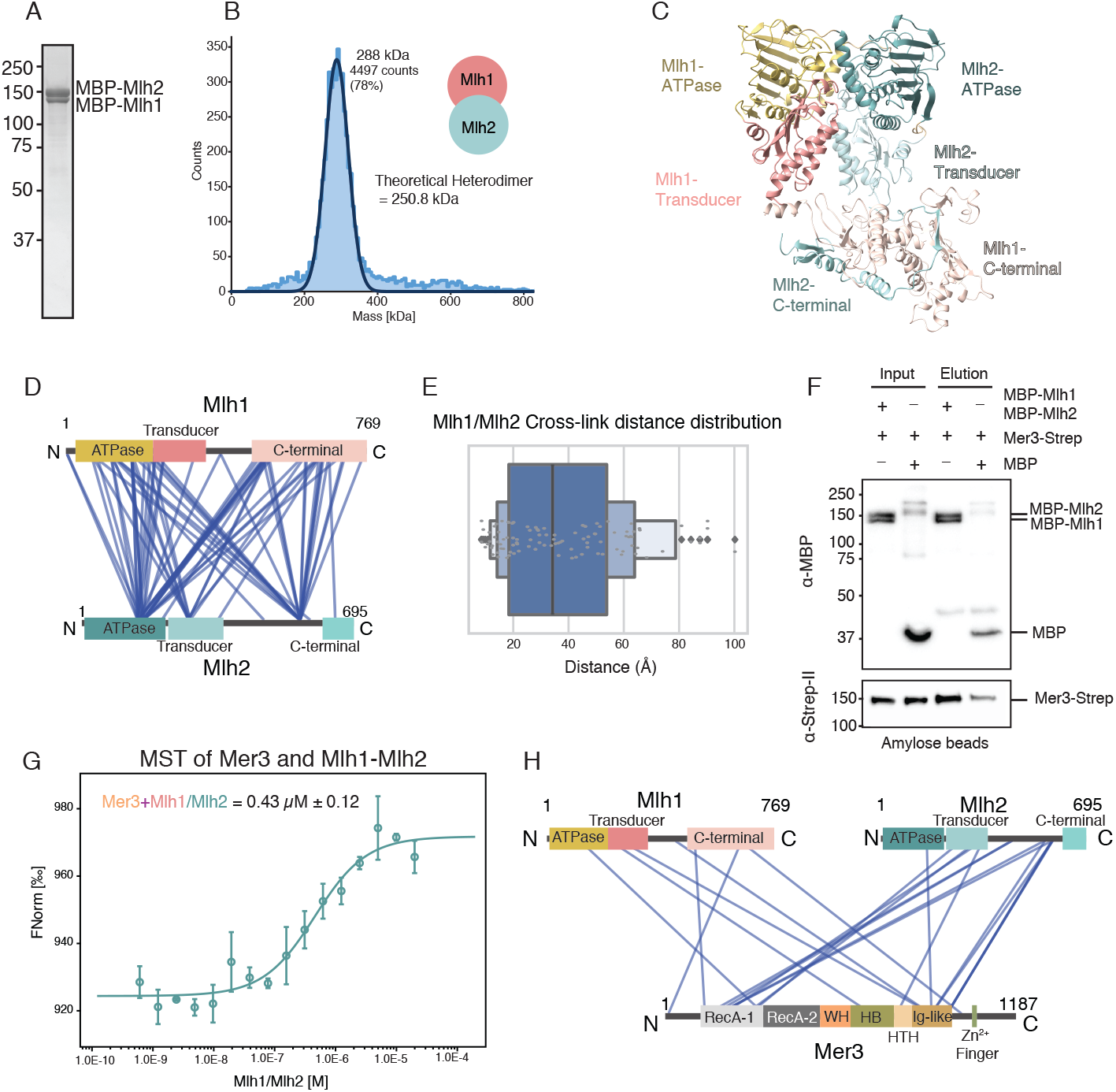
Analysis and quantification of the Mer3/Mlh1/Mlh2 complex. A) Coomassie stained SDS-PAGE showing a representative purification of MBP-Mlh1/MBP-Mlh2 complex (See Supplementary Figure 2A for corresponding chromatograph). B) Mass Photometry profile of purified MBP-Mlh1/MBP-Mlh2 complex (30 mM concentration). We recorded a single species with a molecular mass consistent with a heterodimer of Mlh1/Mlh2. C) AlphaFold2 Multimer (Evans et al., 2021) model of S. cerevisiae Mlh1/Mlh2 heterodimer (see Supplementary Figure 2B and C for quality assessment). Mlh1 is coloured in mustard and pink, and Mlh2 in cyan. Both proteins share the same domain organisation, from N- to C-terminal; ATPase D) domain, Transducer domain, C-terminal domain. E) Visualisation of XL-MS data from purified MBP-Mlh1/MBP-Mlh2 complex, created using XVis (Grimm et al., 2015). Domains are coloured as in C). MBP is removed from the visualisation due to the inability to unambiguously assign cross-links. Intra-chain cross-links are not shown for greater clarity. Only high confidence cross-links (score >50; FDR>1%) are shown. F) Cross-link distance distribution profile from XMAS (Lagerwaard et al., 2022), based on the AF2 model shown in C). Here we indicate that the majority of cross-links are within the accepted distance for the DSBU cross-linker. The exceptions are most likely due to the flexibility of the C-terminal domains of Mh1 and Mlh2 relative to the N-terminus. G) Amylose bead pulldown (binding to MBP), visualised with western blotting (anti-MBP against MBP-Mlh1 and MBP-Mlh2; anti-Strep against Mer3-Strep). We confirm that our recombinant MBP-Mlh1/MBP-Mlh2 complex binds to Mer3 as previously shown (Duroc et al., 2017). H) Microscale thermophoresis (MST) analysis of Mer3 binding to MBP-Mlh1/MBP-Mlh2 complex. Mer3 was fluorescently labelled (see methods) and MBP-Mlh1/MBP-Mlh2 titrated against Mer3. Experiments were carried out in triplicate and the KD of 0.43 M was determined from the fitting curve in the NanoTemper Affinity Analysis v2.3 software (NanoTemper Technologies GmbH). I) Visualisation of XL-MS data of the DSBU cross-linked Mer3/MBP-Mlh1/MBP-Mlh2 complex. MBP is removed from the visualisation due to the inability to unambiguously assign cross-links. Intra-chain cross-links are not shown for greater clarity. Only high confidence cross-links (score >50; FDR>1%) are shown.

The publicly available AlphaFold2 models are currently based on monomeric proteins. To better interpret the results from XL-MS we made an AF2 model of Mlh1/Mlh2 using AlphaFold2 multimer (Evans *et al*., 2021) (Figure 2C). The multimeric model prediction quality was high for the ATPase and transducer regions of both Mlh1 and Mlh2 (Predicted Aligned Error <10 Å, pLDDT > 50) (Supplementary Figure 2B and C). Both Mlh1 and Mlh2 have the same domain organisation; an N-terminal ATPase domain, followed by a transducer domain, an unstructured region, and a C-terminal domain (CTD). The structure of the C-term domain of Mlh1 protein was predicted to be accurate however the general orientation, relative to the ATPase domain and the transducer domain, was low of confidence based on the PAE plot. The C-term domain of Mlh2, as well as the unstructured regions of both proteins, couldn’t be predicted with high confidence.

We characterised the Mlh1/Mlh2 complex using XL-MS and the DSBU crosslinker (Figure 2D). We observed that the majority of the high confidence crosslinks detected between Mlh1 and Mlh2 are broadly distributed on the sequence of the Mlh1 protein whereas they concentrate on two distinctive locations in Mlh2. One of them is in the ATPase domain (K159) and the other one is the N-terminal region of the C-terminal domain (K560) indicating that these two regions of the Mlh2 may be involved in interaction with Mlh1. The N-terminal ATPase-transducer domains of Mlh1 and Mlh2 form a heterodimer that is nearly identical to the N-terminal domain of homodimeric MutL protein (RMSD 1.07Å over 220 amino acids) (Ban *et al*., 1999). We plotted intra- and inter-chain crosslinks on the AF2 Mlh1/Mlh2 model, and found that the overall distribution of cross-link distances is consistent with a generally accurate model (Figure 2E). However a number of long-distance outliers point towards some flexibility within the structure, particularly for the unstructured regions and the C-terminal domain.

In order to further characterise Mer3 and Mlh1/Mlh2 interaction, we first performed an in vitro pull-down assay to confirm that our purified proteins can form a complex (Figure 2F). We characterised the strength of this interaction using microscale thermophoresis (MST) utilising fluorescently labelled Mer3, and determined a KD of 436+/-122 nM (Figure 2G). Next, we determined the structural organisation of the reconstituted Mer3/Mlh1/Mlh2 heterotrimeric complex using XL-MS (Figure 2H). It was previously shown that Mer3 interacted with the Mlh1/Mlh2 complex, at least in part via a conserved region of the Mer3 Ig-like domain (Duroc *et al*., 2017). Consistent with this, we observed several cross-links between the Mer3 Ig-like domain and both Mlh1 and Mlh2. In addition we also observed a large number of cross-links between Mlh2 and RecA-1 of Mer3. We mapped the Mlh1/Mlh2 cross-links onto the surface of the Mer3 AF2 model (Supplementary Figure 2D). One residue (S901) within the Ig-like domain of Mer3 was observed cross-linking to both subunits. Consistent with this observation is that S901 is proximal to R893, which when mutated to glutamic acid, ablates the interaction between Mer3 and Mlh1/Mlh2 (Duroc *et al*., 2017).

### Mer3 interacts with Top3, Rmi1 and the Top3/Rmi1 complex

It was previously inferred that one of the ZMM proteins must antagonise the anti-crossover activity of Sgs1 helicase (Jessop *et al*., 2006). However, recent IP-MS studies had not revealed any potential interaction partners of Mer3 that could facilitate this (Vernekar *et al*., 2021). Given that Sgs1 activity is also required for normal CO formation in meiosis (Oh *et al*., 2007), we speculated any ZMM interaction that could directly or indirectly down-regulate Sgs1 might be transient, and therefore not amenable to proteomics. Thus we made use of a small-scale yeast-two-hybrid screen. We evaluated the interaction of Mer3 with several known components of the cross-over pathway. We found that Mer3 can interact with both Rmi1 and Top3 (Figure 3A). Rmi1 and Top3 are both co-factors for Sgs1, promote its helicase activity ((Cejka *et al*., 2010; Kasaciunaite *et al*., 2019)) and form a so-called “STR complex” (Chang *et al*., 2005). To confirm that Mer3 physically interacts with the Top3/Rmi1 complex in meiosis, we performed a co-immunoprecipitation experiment from *S. cerevisiae* SK1 strain after 6 hours in meiosis. C-terminally tagged Mer3 co-immunoprecipitated with C-terminally HA-tagged Top3 thus confirming that both proteins can associate with one another during meiosis (Figure 3B).

**Figure 3.**
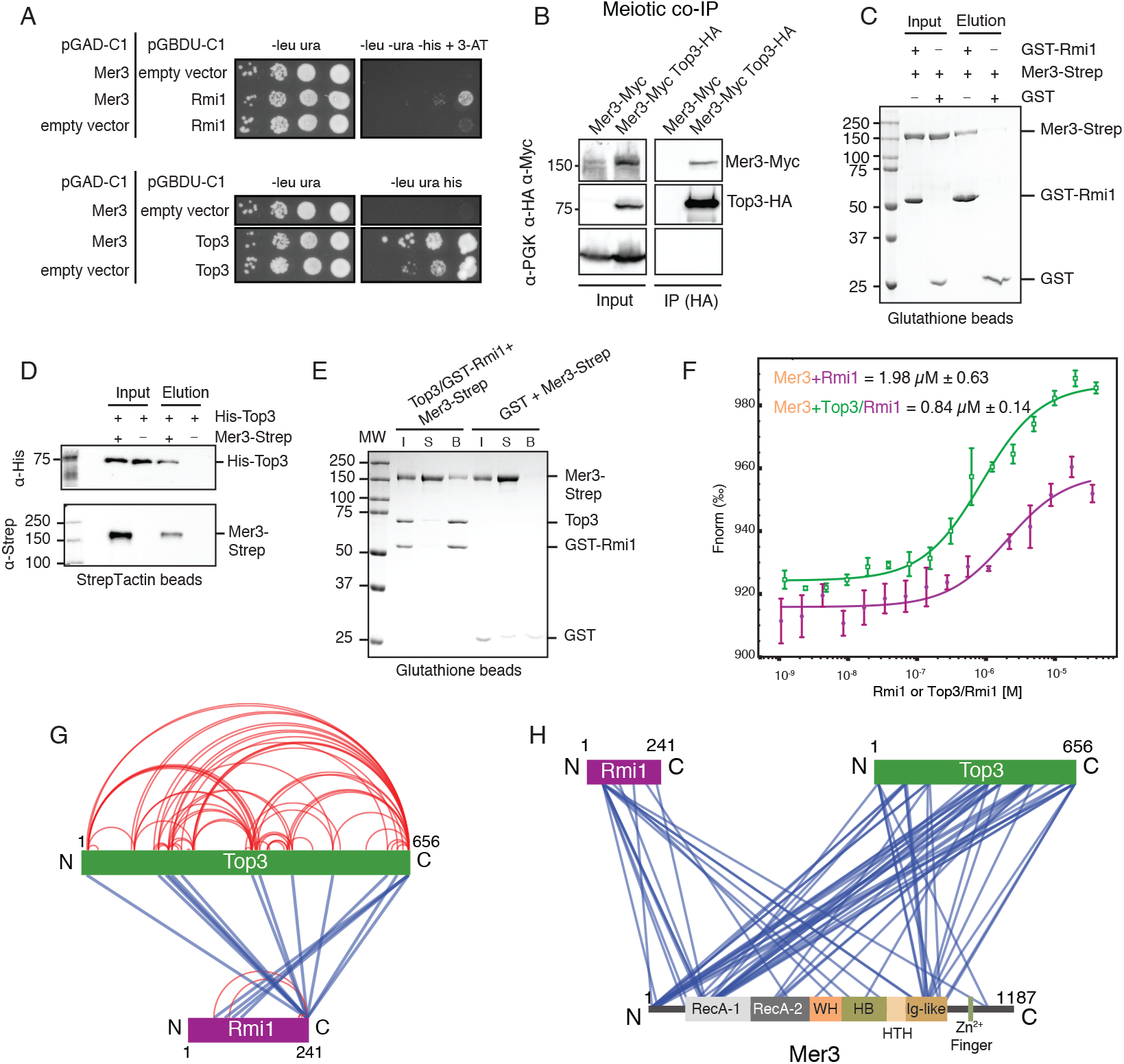
Mer3 binds to Top3 and Rmi1. A) Y2H screen identified a specific interaction between Mer3 and Rmi1 (top panel) and Mer3 and Top3 (bottom panel). B) Meiotic co-IP; after 6 hours in sporulation media lysates were precipitated on anti-HA beads. Anti-Myc western blotting was used to detected Myc-tagged Mer3 in the presence (right lane) or absence (left lane) of HA-tagged Top3. C) Glutathione pulldown of recombinant GST-tagged Rmi1 against Mer3-Strep. GST alone was used as a control for non-specific binding to Mer3. D) StrepTactin pulldown of recombinant Mer3-Strep against His-Top3. Western blotting (anti-Strep and anti-His) was used to confirm the specificity of binding. E) Glutathione pulldown of recombinant GST-tagged Rmi1/Top3 complex against Mer3-Strep. GST alone was used as a control for non-specific binding to Mer3. F) Microscale thermophoresis (MST) of Mer3 binding to Rmi1 (purple) or Top3/Rmi1 complex (green). Mer3 was fluorescently labelled and either Rmi1 alone or Top3/Rmi1 titrated against Mer3. Experiments were carried out in triplicate and the KD of 1.98 M (for Rmi1 alone) or 0.84 M (for Top3/Rmi1) was determined from the fitting curve in the NanoTemper Affinity Analysis v2.3 software (NanoTemper Technologies GmbH). G) Visualisation of Top3/Rmi1 complex XL-MS experiment (using XVis (Grimm et al., 2015)). Intra-chain cross-links are shown in red; inter-chain cross-links in blue. Only high confidence cross-links (score >50; FDR>1]%) are shown. H) Visualisation of Mer3/Top3/Rmi1 complex XL-MS experiment (using XVis (Grimm et al., 2015)). Intra-chain cross-links are not shown for greater clarity. Only high confidence cross-links (score >50; FDR>1%) are shown.

To further study the nature of the Mer3/Top3/Rmi1 interaction we purified both Rmi1 and Top3 as well as a Top3/Rmi1 complex from insect cells (Supplementary Figure 3A). We confirmed the complex formation using these recombinant proteins in an in vitro pulldown assay using C-terminally 2xStrepII-tagged Mer3 (Mer3-Strep) and either GST-Rmi1 or His-Top3. We detected an interaction between both Mer3-Strep and GST-Rmi1 (Figure 3C) and His-Top3 (Figure 3D), indicating that Mer3 makes physical contacts with both proteins. To determine whether both interactions are compatible we also carried out a pulldown using Top3/GST-Rmi1 complex. In these combinations, proteins were also interacting indicating that Mer3 interacts with the Top3/Rmi1 complex (Figure 3E).

We characterised the affinity of the Mer3 to Top3/Rmi1 interaction using microscale thermophoresis (MST). Measured KD of Mer3 binding to Top3/Rmi1 complex was 844+/-148 nM (Figure 3F, green). Importantly, when we measured the binding for Rmi1 alone, the binding was weaker (1.99+/-0.63 μM; Figure 3F, purple) confirming the cooperative nature of the Mer3-Top3/Rmi1 complex assembly. We used AlphaFold2 to predict the structure of the Top3/Rmi1 complex. Overall confidence in the model quality was very high (Supplementary Figure 3B and C) and the model strongly resembles the experimentally determined structure of the human TopoIIIα-RMI1 complex (RMSD of 1.01Å over 444 residues) (Bocquet *et al*., 2014). To evaluate the AF2 model of Top3/Rmi1, we carried out XL-MS on the purified Top3/Rmi1 complex (Figure 3G). 80% of the high confidence crosslinks were consistent with the model (Supplementary Figure 3D).

XL-MS was also used to study the interaction between the Top3/Rmi1 complex and Mer3 (Figure 3H). Top3 showed the most extensive cross-linking with Mer3, with Top3 cross-links clustering in three regions of Mer3; the N-terminal unstructured region, RecA-1 and the Ig-like domain (Figure 3H). We modelled the Top3/Rmi1 cross-links onto the surface of the Mer3 AF2 model (Supplementary Figure 3E). This revealed that the cross-links cluster on one face of the Mer3 molecule made up of the RecA-1 and Ig-like domains, indicating a likely binding site on Mer3.

### Mer3 interaction with Mlh1/2 is compatible with Top3/Rmi1 binding forming a 5-subunit complex

Given that the Ig-like domain of Mer3 is involved in binding to Mlh1/Mlh2 complex (Duroc *et al*., 2017) we tested whether our newly identified binding Top3/Rmi1 by Mer3 is compatible with the Mer3 Mlh1/Mlh2 interaction. We carried out a pulldown using purified Mer3-Strep and Top3/GST-Rmi1 on MBP-Mlh1/MBP-Mlh2 (Figure 4A, lanes 2 and 6). All proteins interacted with each other forming a 5-subunit “supercomplex”. Top3/Rmi1 complex interacted with Mlh1/Mlh2 complex also in the absence of Mer3 (Figure 4A, lanes 3 and 7), as had been previously suggested (Wang and Kung, 2002), and is strongly indicative of a potential cooperative assembly. We again determined the topological structural organisation of the Mer3/Mlh1/Mlh2/Top3/Rmi1 complex using XL-MS (Figure 4B). While the distribution of cross-links between Mer3 and Mlh1/Mlh2 and Top3/Rmi1 were largely similar to the subcomplexes (Figure 2H and 3H) we also found crosslinks between Top3 and Mlh1 as well as between Rmi1 and Mlh2, consistent with our observation that Mlh1/Mlh2 directly interacts with Top3/Rmi1, and thus indicative of a cooperative assembly. We mapped the Mlh1/Mlh2 and Top3/Rmi1 cross-links onto the surface of Mer3 (Figure 4C). We observe that the majority of the cross-links congregate on a single surface made up of RecA-1 and the Ig-like domains. Again this is indicative of a cooperative assembly involving one face of Mer3 that interacts with all four components.

**Figure 4.**
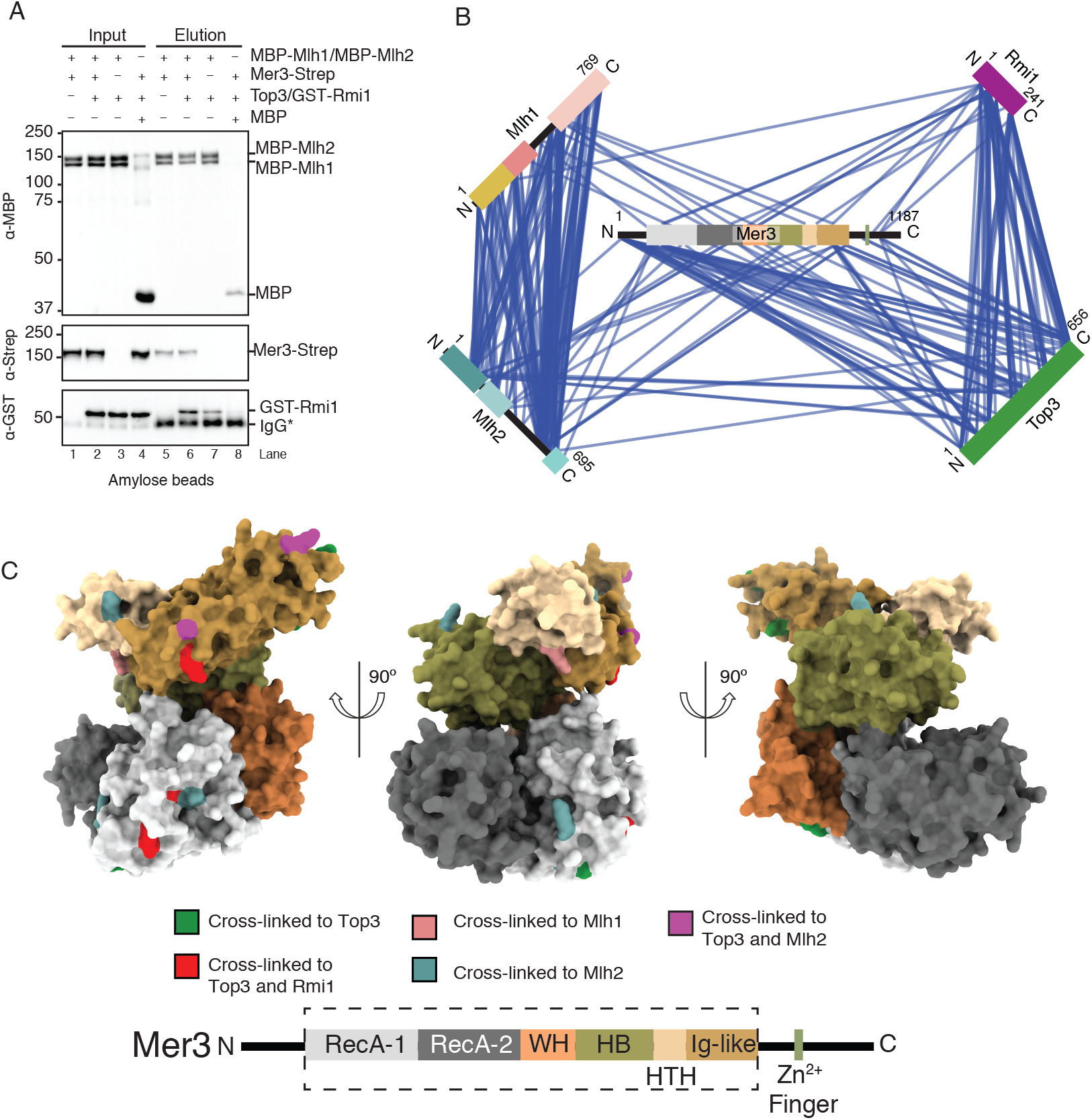
Formation and analysis of a 5-subunit Mer3/Mlh1/Mlh2/Top3/Rmi1 complex. A) Amylose pulldown of recombinant MBP-Mlh1/MBP-Mlh2 complex against Mer3-Strep (lanes 1 and 5) Mer3-Strep and Top3/GST-Rmi1 (lanes 2 and 6); Top3/GST-Rmi1 (lanes 3 and 7). MBP alone serves as a control for background binding (lanes 4 and 8). Western blotting was used to identify the differently tagged components as indicated. B) Visualisation of Mer3/Top3/Rmi1/Mlh1/Mlh2 complex XL-MS experiment (using XVis (Grimm et al., 2015)). Fusion tags and Intra-chain cross-links are not shown for greater clarity. Only high confidence cross-links (score >50; FDR>1C) Surface representation of the structured part of the Mer3 AF2 model. Domains coloured according to the domain cartoon below. Dotted line on domain cartoon represents the structured part of the AF2 prediction, and thus the part of the model shown. Residues of Mer3 that cross-linked to Mlh1, Mlh2, Top3 and Rmi1 are coloured as indicated.

### Mer3 and Sgs1 compete with each other for binding to the Top3/Rmi1 complex

Sgs1 helicase, together with Top3/Rmi1 complex disassemble crossover intermediates and prevent crossover formation. Abolishing the interaction however reduces the activity of both Sgs1 and the Top3/Rmi1 complex (Cejka *et al*., 2010, 2012). Given that both Mer3 and Sgs1 are helicases with some structural similarity we asked whether Mer3 competes with Sgs1 for interaction with Top3/Rmi1 complex. We performed a competitive pulldown where we tested whether increasing amounts of Sgs1 can outcompete Mer3 bound to Top3/Rmi1. In this assay, we used only the N-terminal fragment of Sgs1 (1-605) that is known to interact with Top3/Rmi1 (Bennett *et al*., 2000) due to our difficulty in obtaining suitable stable full-length recombinant Sgs1 (Figure 5A). Triplicates of the same experiment showed a reproducible competition (Figure 5B). We also carried out the same experiment with a shorter fragment of Sgs1 (1-107), containing the previously identified minimum binding region (Bennett *et al*., 2000) which showed a similar effect (Supplementary Figure 4A).

**Figure 5.**
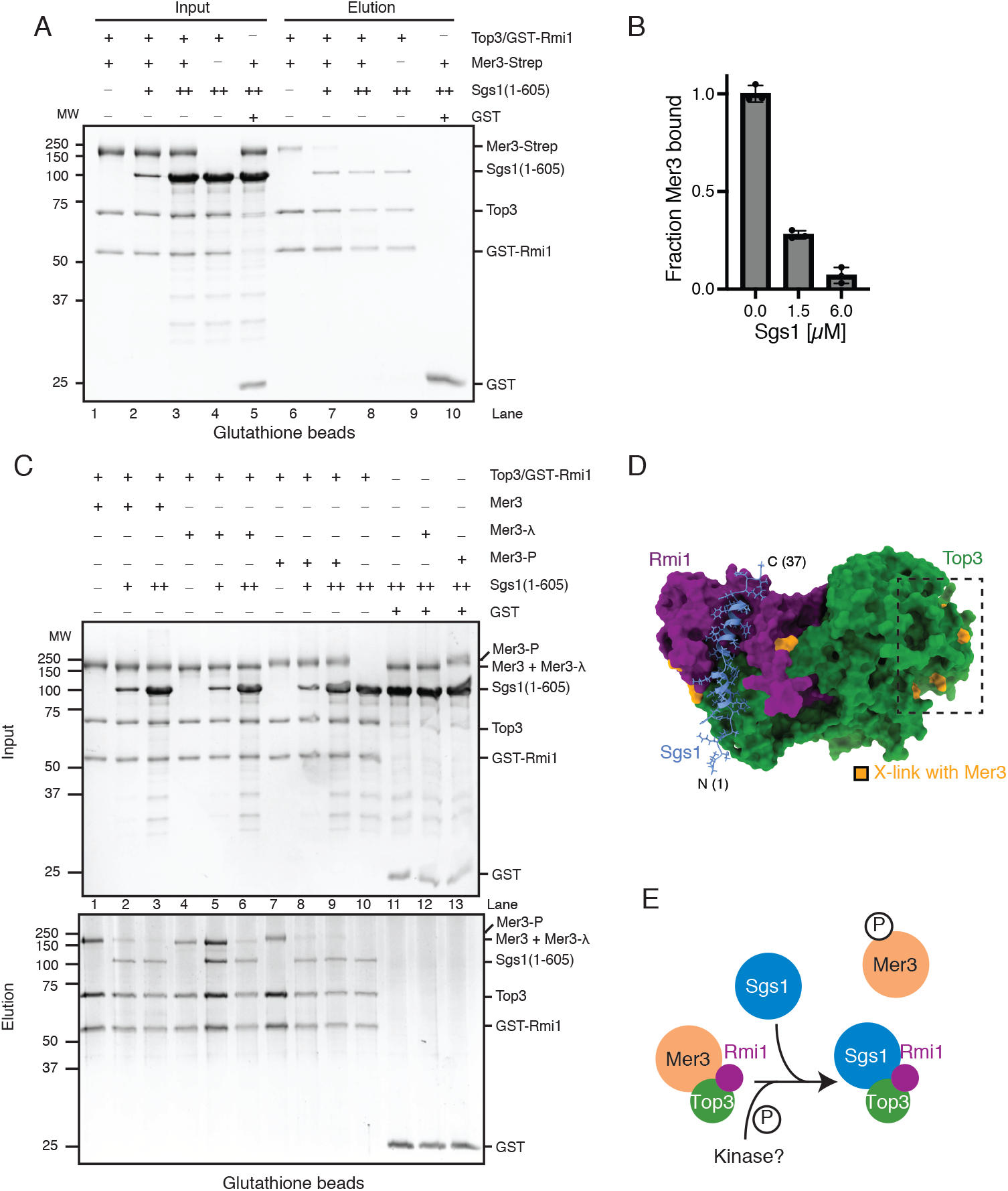
Mer3 and Sgs1 helicases bind competitively for Top3/Rmi1 in a phosphor -dependent manner. A) Glutathione pulldown of recombinant Top3/GST-Rmi1 against Mer3-Strep (lanes 1 and 6), and in increasing concentrations of Sgs11-605 (1.5 M Sgs11-605 lanes 2 and 7; 6 M Sgs11-605 lanes 3 and 8). Lanes 4 and 9 show the binding of Sgs11-605 to Top3/GST-Rmi1 in the absence of Mer3-Strep, and lanes 5 and 10 show the background binding of both Mer3-Strep and Sgs11-605 to GST alone. B) Quantification of lanes 6, 7 and 8 from A). Circles are measurements from three independent experiments, error bars show the SD of these three measurements. C) Glutathione pulldown, carried out in an equivalent manner to A), but using three different versions of Mer3-Strep. Mer3, purified as previously; Mer3-^λ^, treated with ^λ^-phosphatase; Mer3-P, purified from cells treated with okadaic acid. D) AF2 structure of the Top3/Rmi1/Sgs1N complex. Surface representation of Top3 (green) and Rmi1 (purple) and a cartoon representation of Sgs1 (blue). The N-terminal helix of Sgs1 (residues 9-34) is predicted to bind to a cleft formed by both Top3 and Rmi1. Residues of Mer3 that cross-link to Top3 and Rmi1 are coloured orange. E) Unphosphorylated Mer3 forms a complex with Top3-Rmi1. Upon phosphorylation by an unknown kinase Mer3 binds more weakly to Top3-Rmi1 allowing displacement by Sgs1 kinase.

Given that Sgs1 was able to displace Mer3 from Top3/Rmi1 at relatively low concentrations, we asked whether the interaction might be modulated through post-translational modifications. We took advantage of the insect cell expression of Mer3 to create three variants. Hyperphosphorylated Mer3 (Mer3-P), was purified from insect cells treated with okadaic acid; “normal” phosphorylated Mer3 was purified from untreated insect cells; dephosphorylated Mer3 (Mer3-λ), was purified from untreated insect cells and subject to lambda-phosphatase treatment. In our pulldown experiment Mer3-λ showed far more resistance to Sgs1 competition, whereas Mer3-P could be more easily displaced from Top3/Rmi1 (Figure 5C).

Our attempts to create an AlphaFold2 model of Mer3/Top3/Rmi1 were not successful, perhaps due to the complex binding mode of Mer3 to Top3/Rmi1 (see discussion). We were however able to create a high-confidence prediction of an N-terminal fragment of Sgs1 bound to Top3/Rmi1 (Supplementary Figure 4B). When we mapped the residues of Top3/Rmi1 that cross-link to Mer3 we found that these residues form two clusters, one on the N-terminus of Top3, and the second around the predicted Sgs1 binding site (Figure 5D). Taken together we show that Mer3 and Sgs1 appear to compete for the same binding site on the Top3/Rmi1 complex (Figure 5E).

### Mer3 inhibits D-loop disassembly mediated by the Sgs1/Top3/Rmi1 complex

Given that Mer3 potentially affects the activity of the STR complex, we reconstituted D-loop formation using yeast meiosis-specific recombinase Dmc1 and RPA protein (Figure 6A). Radioactively labelled ssDNA (90-mer) was first incubated with Dmc1 recombinase to form a presynaptic filament. After short incubation with RPA, D-loop formation was initiated by addition of supercoiled plasmid DNA. Then, STR complex (15 nM) was added to the indicated reactions which resulted in robust disruption of the D-loop after 10 min of incubation (30% of relative D-loop yield formed in the absence of STR) (Figure 6B and C). Interestingly, increasing concentrations of Mer3 were able to inhibit the D-loop disassembly by STR complex. 20-fold excess of Mer3 (300 nM) over Sgs1 resulted in a 70% relative yield of D-loop (compared to 30% in the absence of Mer3). Our model makes the assumption that it is the protein binding activity of Mer3, rather than the helicase activity, that inhibits the STR complex. To test this, we made use of an ATP hydrolysis deficient mutant of Mer3, Mer3 K167A. Quantification of triplicate D-loop assays revealed that Mer3K167A showed the same inhibition of STR complex as wild type Mer3 (Figure 6 D and E).

**Figure 6.**
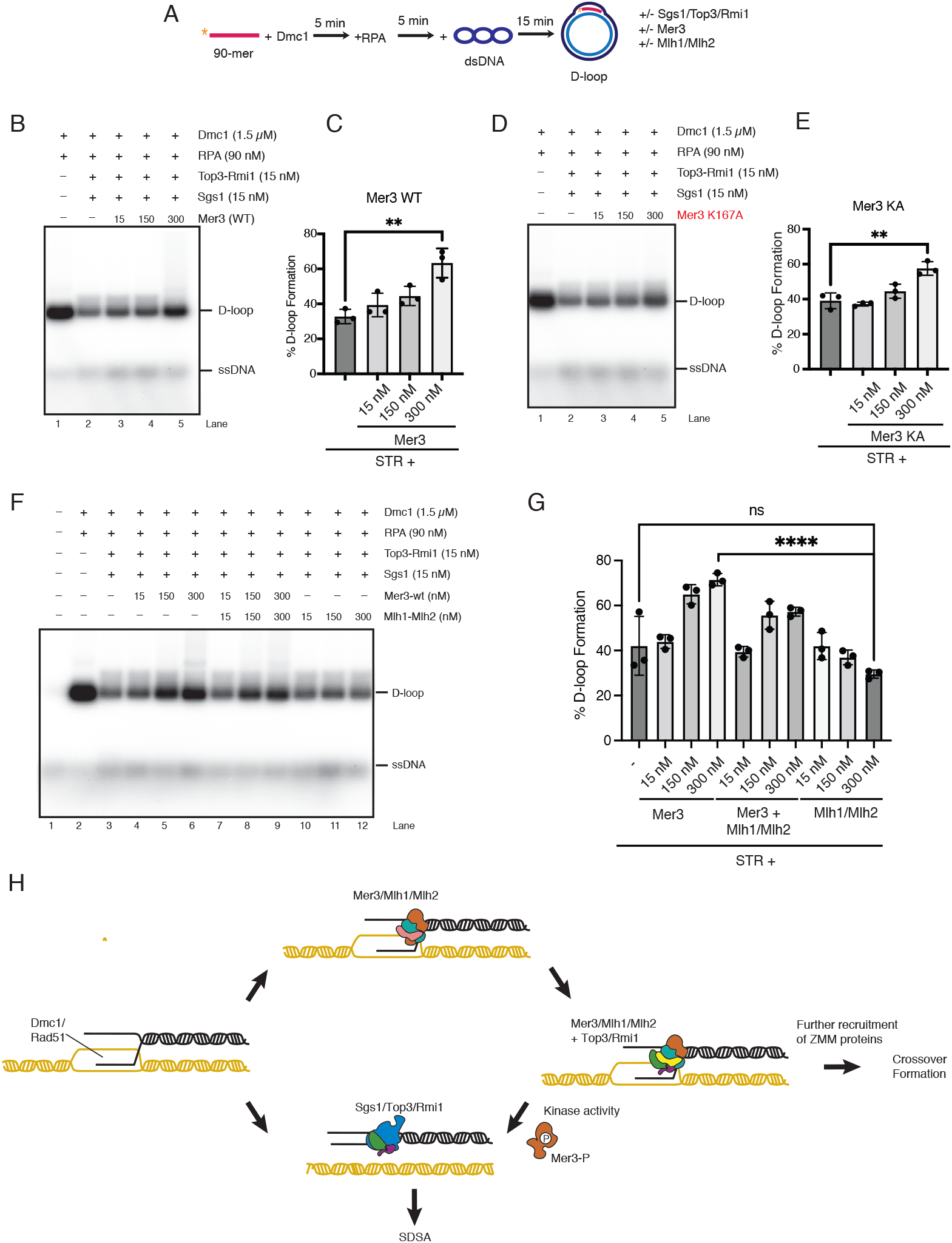
Mer3 protects D-loops from disassembly by STR complex. A) Schematic of the D-loop disassembly experiment. 5 radio-labelled 90-mer ssDNA was incubated in the presence of Dmc1, followed by RPA, followed by a 15-minute incubation with circular dsDNA to form initial “D-loop” like DNA repair intermediates. These D-loops were then further incubated for 10 minutes in the presence of STR complex, or STR complex with Mer3 or Mlh1/Mlh2. B) Image of samples run on a 0.9% agarose gel. Lane 1 shows the formation of D-loops in the absence of STR or Mer3. Lane 2 shows the effect of the STR complex on D-loops. Lanes 2-5 shows the effect of increasing wild-type Mer3 concentration on the formation of D-loops. C) Quantification of experiments as shown in B). The formation of D-loops in the absence of STR was taken to be 100% formation. Solid circles show values measured from three independent experiments. Error bars show the SD from these three experiments. An unpaired t-test was used to determine statistical significance of the experiments in lanes 2 and 5. D) Image of samples run on a 0.9% agarose gel. Lane 1 shows the formation of D-loops in the absence of STR or Mer3KA. Lane 2 shows the effect of STR complex on D-loops. Lanes 2-5 shows the effect of increasing ATPase dead Mer3 concentration on the formation of D-loops. E) Quantification of experiments as shown in D). The formation of D-loops in the absence of STR was taken to be 100% formation. Solid circles show values measured from three independent experiments. Error bars show the SD from these three experiments. An unpaired t-test was used to determine statistical significance of the experiments in lanes 2 and 5. F) Image of samples run on a 0.9% agarose gel. Lane 1 shows DNA alone. Lane 2 the formation of D-loops in the absence of STR, Mer3 or Mlh1/Mlh2. Lane 3 shows the effect of the STR complex on D-loops. Lanes 4-6 show the effect of increasing wild-type Mer3 concentration on the formation of D-loops. Lanes 7-9 show the effect of increasing wild-type Mer3 and Mlh1/Mlh2 complex concentration on the formation of D-loops. Lanes 10-12 show the effect of increasing Mlh1/Mlh2 complex concentration on the formation of D-loops. G) Quantification of experiments as shown in F). The formation of D-loops in the absence of STR was taken to be 100% formation. Solid circles show values measured from three independent experiments. Error bars show the SD from these three experiments. An unpaired t-test was used to determine statistical significance of the experiments in lanes 3 and 12 and lanes 6 to 12. H) Cartoon of the proposed mechanism of Mer3 mediated D-loop protection during meiosis. Mer3 captures initial D-loops cooperatively with Mlh1/Mlh2. Uncaptured D-loops are disassembled by STR and repaired via the SDSA pathway. Mer3 recruits Top3-Rmi1 at D-loops, thus protecting these D-loops from disassembly by STR complex. Mer3 binding to Top3/Rmi1 is regulated by an unknown kinase that may allow some initially captured D-loops to be subsequently disassembled by STR.

Since Mlh1/Mlh2 potentially contributes to a cooperative assembly with Mer3 and Top3/Rmi1 we asked whether the addition of Mlh1/Mlh2 influences the D-loop protection activity of Mer3. Addition of Mlh1/Mlh2 to the Mer3-STR assay did not improve Mer3 mediated D-loop protection, indeed there was a mild reduction in the Mer3 mediated protection (Figure 6F and G). Since Mer3 is a tight binder of D-loops (Duroc *et al*., 2017) (Supplementary Figure 1A), we asked whether Mer3 might antagonise STR activity simply by binding to D-loops. To test this, we analysed the effect of Mlh1/Mlh2 alone on STR mediated D-loop disassembly. Mlh1/Mlh2 shows binding to D-loops with a similar affinity as Mer3 (Duroc *et al*., 2017). If the Mer3-effect is simply D-loop binding, we would expect to see an effect by titrating Mlh1/Mlh2 against the STR complex. Interestingly, we saw no effect on STR mediated D-loop disassembly with the addition of Mlh1/Mlh2 complex (Figure 6F and G). We therefore suggest that Mer3 inhibits the D-loop disassembly activity of the STR complex through its binding to Top3/Rmi1.

## Discussion

Mer3 is a highly conserved member of the ZMM group of proteins that promotes meiotic crossover formation. No experimental structure of Mer3 or its functional orthologs have been described to date. Recent breakthroughs in protein structure prediction, combined with previous biochemical work on both Mer3 and other similar helicases has allowed us to undertake a basic structure-function characterisation of Mer3. It should be noted however the AF2 predictions of Mer3 appear to be at least superficially similar to Mer3 models generated using homology modelling (Duroc *et al*., 2017). More detailed structure-function analysis was compounded by the observation that Mer3 can form homodimers.

Biophysical analysis of purified Mer3 shows dimer formation at high concentrations, and monomers at low concentrations, and that this dimerisation depends upon the N- and C-terminal unstructured regions. We propose that in the meiotic nucleoplasm Mer3 is monomeric, but that once D-loops start to form through the course of meiotic prophase, Mer3 forms dimers on D-loops (likely aided by binding to Mlh1/Mlh2, see below). Currently we have no structural information on the organisation of the dimer, beyond the three identified “self” cross-links from XL-MS (Figure 1F). One interesting possibility, however, is derived from the spliceosomal Brr2 helicase, to which Mer3 is clearly structurally related. Brr2 contains two helicase cassettes in the polypeptide, with the C-terminal cassette lacking catalytic activity, but instead mediating protein-protein interactions. As such we might speculate that one Mer3 subunit is in contact with the D-loop, and extends the D-loop as previously suggested (Mazina *et al*., 2004), and the second mediates protein-protein interactions.

One previously identified protein interaction of Mer3 is with the MutLβ complex Mlh1/Mlh2 (Duroc *et al*., 2017). We characterised this interaction further, finding a submicromolar KD, and identifying candidate contact regions and residues through XL-MS. Our data is entirely consistent with the previous work from the Borde laboratory on Mer3 and Mlh1/Mlh2, which showed a role of the Ig-like domain of Mer3 in binding to Mlh1/Mlh2. In addition we identify the 1st RecA-like domain of Mer3 as a second potential binding interface with Mlh1/Mlh2. A new protein-protein interaction that we identify is between Mer3 and Top3/Rmi1. We measured the binding affinity to be 2-fold less than Mer3 to Mlh1/Mlh2, which might partly explain why this interaction was not previously identified with proteomic approaches in both directions (Vernekar *et al*., 2021; Wild *et al*., 2019). We also found that, while the interactions with Top3/Rmi1 and Mlh1/Mlh2 appear to utilise the same surface of Mer3, they are compatible (and likely cooperative) with one another. As such we speculate that Mer3 and Mlh1/Mlh2 cooperate in the context of meiotic D-loops to bind to Top3/Rmi1.

What is the function of Mer3 binding to Top3/Rmi1? We found that Mer3 binds competitively with the Sgs1 helicase (a functional ortholog of BLM helicase) to Top3/Rmi1, disrupting the formation of the STR complex. By combining XL-MS data with an AF2 model of the Sgs1/Top3/Rmi1 structure, we suggest that Mer3 binds to the same cleft on Top3/Rmi1 that is occupied by an N-terminal alpha-helix of Sgs1. By making use of differently phosphorylated versions of Mer3, we find that the higher the level of phosphorylation of Mer3, the more easily Sgs1 can displace Mer3 from Top3/Rmi1. This hints at a mechanism by which an increase in cellular kinase activity serves to disrupt the Mer3/Top3/Rmi1 complex, and potentially allow Sgs1 to bind (Figure 6H). This regulated interaction is consistent with the observation that while Sgs1 is an “anti-crossover” helicase, its functionality is also required for normal crossover formation. Therefore we suggest that the Mer3 binding to Top3/Rmi1 is transient, and possibly only at a limited number of loci within the genome.

In a D-loop disassembly assay we find that titrating Mer3 against STR complex reduces the level of D-loop disassembly; presumably by physically disrupting the STR complex. Two controls give further weight to this argument. Firstly, the helicase activity of Mer3 is not required for D-loop “protection” and secondly other tight D-loop binders, in this case Mlh1/Mlh2 complex, do not protect D-loops from STR complex mediated disassembly. One surprising observation is that the combination of Mlh1/Mlh2 with Mer3 does not provide additional protection to D-loops, indeed the addition of Mlh1/Mlh2 seems to partly mitigate the Mer3 mediated protection. We suggest that this might be due to an excess of Mlh1/Mlh2 outcompeting Mer3 from D-loops. Furthermore this mild abrogation of Mer3 mediated protection in a bulk assay might hide the fact that, for example, the residence time of Mer3 on D-loops might be increased in the presence of Mlh1/Mlh2, due to a change in kinetics.

Here we show for the first time a physical connection between the ZMM proteins and the STR complex. We suggest that Mer3 replaces Sgs1 as the helicase in the STR complex, which has the opposite effect of stabilising rather than disassembling D-loops. We suggest that this is cooperative with other factors, in particular Mlh1/Mlh2, but that this may also be cooperative with other ZMM factors. Indeed our work suggests that Mer3 would be one of the first ZMM proteins that binds at SEI intermediates, which would be in line with what has been shown previously (Duroc *et al*., 2017; Pyatnitskaya *et al*., 2019).

Several outstanding questions remain to be answered by future experiments. Firstly, when bound to Top3/Rmi1, what interplay is there between the enzymatic activities, i.e. the decatenase and helicase centres? Secondly, besides Mlh1/Mlh2, what other protein factors influence the formation of the Mer3/Top3/Rmi1 complex and potentially contribute to D-loop protection? Thirdly, which kinase is regulating the interaction between Mer3 and Top3/Rmi1, and is there a synergistic effect with phosphorylation of other components in the system (e.g. Sgs1 (Grigaitis *et al*., 2020)). Finally, given the level of conservation of all of the components discussed here, is the interaction between Mer3 and Top3/Rmi1 conserved in other species? Future work, particularly with larger reconstitutions, utilising high-resolution structural biology and single molecule approaches will aim to address these questions.

## Materials and Methods

### Cloning

Sequences of Saccharomyces cerevisiae MER3, MLH1, MLH2, TOP3, RMI1, and SGS1 were derived from SK1 strain genomic DNA. Due to the presence of an intron in MER3, this was amplified as two separate fragments and Gibson assembled. Plasmids used for protein expression were cloned as previously described from the InteBac (Altmannova *et al*., 2021) and biGBac vector suites (Weissmann *et al*., 2016). Baculovirus was generated via the EMBacY cells, part of the MultiBac expression system (Bieniossek *et al*., 2012).

### Protein Expression

Mer3, Mlh1, Mlh2, Top3, Rmi1, and full-length Sgs1 were produced in the Hi-5 cell line derived from the cabbage looper Trichoplusiani. Bacmids for expression were produced in EmBacY cells and subsequently used to transfect Sf9 cells to produce baculovirus. Amplified baculovirus was used to infect Hi-5 cells in 1:100 dilution prior to 72-hour cultivation and harvest. Cells were washed twice with 1xPBS, frozen in liquid nitrogen and the pellets were stored at −80°C.

### Protein purification

#### Mer3

Mer3 was produced as a C-terminal Twin-StrepII tag in Hi5 insect cells using the same expression conditions as described above.To purify Mer3-Strep, cells were resuspended in lysis buffer (50 mM HEPES pH 6.8, 300 mM NaCl, 5% glycerol, 0.1% Triton-X 100, 1 mM MgCl2, 5 mM β-mercaptoethanol). Resuspended cells were lysed using an EmulsiFlex C3 (Avestin) in presence of Serva Protease-Inhibitor Mix and Benzonase before clearance at 130,000g at 4°C for 1 hour. Cleared lysate was applied on a 5 mL Strep-TactinXT column (iba) and extensively washed with lysis buffer. Mer3 constructs were eluted with a lysis buffer containing 50 mM Biotin. Eluted protein was passed through a HiTrap Heparin HP affinity column (Cytiva) pre equilibrated with the loading buffer (50 mM HEPES pH 6.8, 300 mM NaCl, 5% glycerol, 1 mM MgCl2, 5 mM β-mercaptoethanol). The proteins were eluted by increasing salt gradient to 1M NaCl. Protein-containing elution fractions were concentrated on Vivaspin 15R, 30,000 MWCO Hydrosart concentrators. The concentrated eluent was loaded on a Superdex 200 16/600 pre-equilibrated in SEC buffer (30 mM MES pH 6.5, 300 mM NaCl, 5% glycerol, 1 mM MgCl2, 1 mM TCEP). Purified protein was concentrated using Vivaspin 15R, 30,000 MWCO Hydrosart concentrators. Dephosphorylated Mer3 (denoted as “Mer3-λ”) was prepared using a similar procedure which included lambda-phosphatase (New England Biolabs, P0753) treatment of concentrated Mer3 before loading on the size-exclusion column. To prepare phosphorylated Mer3 (denoted as “Mer3-P”), 100 nM okadaic acid was added to the Hi5 cells for the last 3 hours before harvesting. Protein was then purified using the same protocol as described above.

#### Mlh1/Mh2

To purify the Mlh1-Mlh2 complex, N-terminal 6xHis-MBP tagged Mlh1 and Mlh2 were cloned into the pBIG1a vector and expressed in Hi5 insect cells using the same expression conditions as described above. The cell pellet was resuspended in lysis buffer (50 mM HEPES pH 6.8, 300 mM NaCl, 5% glycerol, 0.1% Triton X-100, 1 mM MgCl2, 5 mM β-mercaptoethanol). Resuspended cells were lysed using an EmulsiFlex C3 (Avestin) in presence of Serva Protease-Inhibitor Mix and Benzonase before clearance at 130,000g at 4°C for 1 hour. Cleared lysate was applied on a 5 mL MBP-trap column (Cytiva) and extensively washed with lysis buffer. Mlh1/Mlh2 complex was eluted with a lysis buffer containing 1 mM maltose. Eluted protein was passed through a HiTrap Heparin HP affinity column (Cytiva) pre-equilibrated with the loading buffer (50 mM HEPES pH 6.8, 300 mM NaCl, 5% glycerol, 1 mM MgCl2, 5 mM β-mercaptoethanol). The proteins were eluted by increasing salt gradient to 1 M NaCl. Protein-containing elution fractions were concentrated on Vivaspin 15R, 30,000 MWCO Hydrosart concentrators. The concentrated eluent was loaded on a Superdex 200 16/600 pre-equilibrated in SEC buffer (30 mM HEPES 6.8, 300 mM NaCl, 5% glycerol, 1 mM MgCl2, 1 mM TCEP). Purified protein was concentrated using Vivaspin 15R, 30,000 MWCO Hydrosart concentrators.

To purify Mlh1-Mlh2 complex (used in D-loop assays), N-terminal 6xHis tagged-Mlh1 and N-terminal Twin-StrepII tagged Mlh2 were cloned into the pBIG1a vector and expressed in Hi5 insect cells using the same expression conditions as described above. The cell pellet was resuspended in the lysis buffer (50 mM HEPES pH 7.5, 300 Mm NaCl, 5% glycerol, 0.01% NP40, 5 mM β-mercaptoethanol, AEBSF, Serva protease inhibitor cocktail). Resuspended cells were lysed by sonication before clearance at 35,000 rpm at 4°C for 1 hour. Cleared lysate was loaded on a 5 mL Strep-Tactin XT 4Flow column (IBA) followed by the first wash using 25 mL of H buffer (20 mM HEPES pH 7.5, 5% glycerol, 0.01% NP40, 1 mM β-mercaptoethanol) containing 500 mM NaCl and the second wash with H buffer containing 150 mM NaCl. Mlh1-Mlh2 complex was eluted with a 50-mL gradient of H buffer containing 150 mM NaCl and 50 mM biotin. Partially purified protein was further loaded onto a 5 mL HiTrap Heparin HP affinity column (Cytiva) pre-equilibrated in H buffer containing 150 mM NaCl and eluted with an increasing salt gradient to 1 M NaCl. The fractions containing Mlh1-Mlh2 complex were then concentrated on a 50 kDa MWCO Amicon concentrator and applied onto a Superose 6 10/300 column (Cytiva) pre-equilibrated in SEC buffer (20 mM HEPES pH 7.5, 300 mM NaCl, 5% glycerol, 1 mM β-mercaptoethanol, 1 mM TCEP). The fractions containing Mlh1-Mlh2 were concentrated on a 50 kDa MWCO Amicon concentrator and stored at −80°C in small aliquots.

#### Top3

Top3 was produced as an N-terminal 6xHis tag in Hi5 insect cells using the same expression conditions as described above. The cell pellet was resuspended in the lysis buffer (50 mM HEPES pH 7.5, 300 mM NaCl, 5% glycerol, 0.01% NP40, 5 mM β-mercaptoethanol, AEBSF). Resuspended cells were lysed by sonication before clearance at 35,000 rpm at 4°C for 1 hour. Cleared lysate was loaded on a 5 mL HiTrap TALON Crude column (Cytiva) followed by a wash using 25 mL of H buffer (20 mM HEPES pH 7.5, 5% glycerol, 0.01% NP40, 1 mM β-mercaptoethanol) containing 150 mM NaCl. Top3 protein was eluted with a 50-mL gradient of 0-450 mM imidazole in an H buffer containing 150 mM NaCl. Partially purified protein was further loaded onto a 5 mL HiTrap Heparin HP affinity column (Cytiva) pre-equilibrated in H buffer containing 150 mM NaCl and eluted with an increasing salt gradient to 1 M NaCl. The fractions containing Top3 protein were then loaded onto a 6 mL ResourceS column (Cytiva) pre-equilibrated in an H buffer containing 150 mM NaCl. and eluted with increasing salt gradient to 1 M NaCl. The peak fractions were concentrated on a 30 kDa MWCO Amicon concentrator and applied onto a Superose 6 10/300 column (Cytiva) pre-equilibrated in SEC buffer (20 mM HEPES pH 7.5, 300 mM NaCl, 5% glycerol, 1 mM β-mercaptoethanol, 1 mM TCEP). The fractions containing Top3 were concentrated on a 30 kDa MWCO Amicon concentrator and stored at −80°C in small aliquots.

#### Top3/Rmi1

To purify the Top3/Rmi1 complex, untagged Top3 and N-terminal GST-tagged Rmi1 were cloned into the pBIG1a vector and expressed in Hi5 insect cells using the same expression conditions as described above. The cell pellet was resuspended in the lysis buffer (50 mM HEPES pH 7.5, 300 mM NaCl, 5% glycerol, 0.01% NP40, 5 mM β-mercaptoethanol, AEBSF). Resuspended cells were lysed by sonication before clearance at 35,000 rpm at 4°C for 1 hour. Cleared lysate was loaded on a 5 mL GSTrap column (Cytiva) followed by wash using 25 mL of H buffer (20 mM HEPES pH 7.5, 5% glycerol, 0.01% NP40, 1 mM β-mercaptoethanol) containing 300 mM NaCl. The Top3/Rmi1 complex was eluted with 50 mL of H buffer containing 100 mM NaCl and 100 mM glutathione. Partially purified protein was further loaded onto a 6 mL ResourceS column (Cytiva) pre-equilibrated in H buffer containing 100 mM NaCl and eluted with an increasing salt gradient to 800 mM NaCl. The peak fractions were concentrated on a 30 kDa MWCO Amicon concentrator and applied onto a Superose 6 10/300 column (Cytiva) pre-equilibrated in SEC buffer (20 mM HEPES pH 7.5, 300 mM NaCl, 5% glycerol, 1 mM β-mercaptoethanol, 1 mM TCEP). The fractions containing the Top3/Rmi1 complex were concentrated on a 30 kDa MWCO Amicon concentrator and stored at −80°C in small aan liquots. To obtain untagged Top3/Rmi1 complex, the concentrated elute fractions from ResourceS column were mixed with 3C HRV protease in a molar ratio of 50:1 and incubated overnight at 4°C. Afterward, the cleaved protein was loaded onto a Superdex 200 10/300 column (Cytiva) with its outlet connected to a 5 mL GSTrap column (Cytiva) pre-equilibrated in SEC buffer (20 mM HEPES pH 7.5, 300 mM NaCl, 5% glycerol, 1 mM β-mercaptoethanol, 1 mM TCEP). The fractions containing untagged Top3/Rmi1 were concentrated on a 10 kDa MWCO Amicon concentrator and stored at −80°C in small aliquots.

#### Sgs1

Sgs1 containing an N-terminal 6xHis-MBP tag and C-terminal 6xHis tag was produced in Hi5 cells using the similar expression conditions as described above with a minor change in using 1:300 dilution of baculovirus. The cell pellet (17 g) was resuspended in the lysis buffer (50 mM HEPES pH 7.5, 300 mM NaCl, 5% glycerol, 0.01% NP40, 5 mM β-mercaptoethanol, AEBSF, Serva protease inhibitor cocktail, and 1 mM PMSF). Resuspended cells were lysed by sonication before clearance at 35,000 rpm at 4°C for 1 hour. Cleared lysate was loaded on a 5 mL MBPTrap column (Cytiva) followed by wash using 35 mL of H buffer (20 mM HEPES pH 7.5, 5% glycerol, 0.01% NP40, 1 mM β-mercaptoethanol) containing 300 mM NaCl. The Sgs1 protein was eluted with a 50-mL gradient of 0-20 mM maltose of H buffer containing 150 mM NaCl. To cleave off N-terminal 6xHis-MBP tag, partially purified protein was mixed with 100 μL 3C HRV protease (6 μg/μL) and incubated overnight at 4°C. Afterward, the cleaved protein was loaded onto a 5 mL HiTrap Heparin HP affinity column (Cytiva) pre-equilibrated in H buffer containing 100 mM NaCl and eluted with an increasing salt gradient from 300 mM to 1 M NaCl. The fractions containing Sgs1 protein were concentrated on a 100 kDa MWCO Amicon concentrator and stored at −80°C in small aliquots. Sgs1(1-107) fragment containing N-terminal 6xHis tag was produced in E. coli strain BL21 STAR. Protein expression was induced by 0.5 mM IPTG at 37°C for 3 hours in TB media supplemented with ampicillin (100 μg/mL). The cell pellet was resuspended in the lysis buffer (50 mM HEPES pH 7.5, 300 mM NaCl, 5% glycerol, 0.01% NP40, 5 mM β-mercaptoethanol, AEBSF, Serva protease inhibitor cocktail). Resuspended cells were lysed by sonication before clearance at 35,000 rpm at 4°C for 1 hour. Cleared lysate was incubated with 800 μL of Ni-NTA agarose (Qiagen) for 1 hour at 4°C. The beads were washed with 20 mL of H buffer containing 150 mM NaCl followed by another wash with 20 mL of H buffer containing 150 mM NaCl and 10 mM imidazole. The protein was eluted in steps with 50, 150, 300, and 500 mM imidazole in H buffer containing 150 mM NaCl. Partially purified protein was further loaded onto a 6 mL ResourceQ column (Cytiva) pre-equilibrated in H buffer containing 150 mM NaCl and eluted with increasing salt gradient to 1 M NaCl. The fractions containing Sgs1 fragment were concentrated on a 10 kDa MWCO Amicon concentrator and applied onto a Superdex 200 10/300 column (Cytiva) pre-equilibrated in SEC buffer (20 mM HEPES pH 7.5, 300 mM NaCl, 5% glycerol, 1 mM β-mercaptoethanol, 1 mM TCEP). The fractions containing Sgs1 protein were concentrated on a 10 kDa MWCO Amicon concentrator and stored at −80°C in small aliquots.

Sgs1(1-605) fragment containing N-terminal 6xHis-MBP tag was produced in E. coli strain BL21 STAR. Protein expression was induced by 0.2 mM IPTG at 18°C overnight in TB media supplemented with ampicillin (100 μg/mL). The cell pellet was resuspended in the lysis buffer (50 mM HEPES pH 7.5, 300 mM NaCl, 5% glycerol, 0.01% NP40, 5 mM β-mercaptoethanol, AEBSF, Serva protease inhibitor cocktail). Resuspended cells were lysed by sonication before clearance at 35,000 rpm at 4°C for 1 hour. Cleared lysate was loaded on a 5 mL MBPtrap column (Cytiva) followed by first wash using 25 mL of H buffer (20 mM HEPES pH 7.5, 5% glycerol, 0.01% NP40, 1 mM β-mercaptoethanol) containing 500 mM NaCl and the second wash using H buffer containing 150 mM NaCl. The protein was eluted with a 50 mL of H buffer containing 150 mM NaCl and 20 mM maltose. Partially purified protein was further loaded onto a 6 mL ResourceQ column (Cytiva) pre-equilibrated in H buffer containing 150 mM NaCl and eluted with increasing salt gradient to 1 M NaCl. The fractions containing Sgs1 fragment were concentrated on a 30 kDa MWCO Pierce concentrator and incubated with 3C HRV protease in a molar ratio of 50:1 overnight at 4°C. Afterwards, the cleaved protein was applied onto a Superose 6 10/300 column (Cytiva) with its outlet connected to a 5 mL GSTrap column (Cytiva) followed by a 5 mL MBPtrap column (Cytiva). All columns were pre-equilibrated in SEC buffer (20 mM HEPES pH 7.5, 300 mM NaCl, 5% glycerol, 1 mM β-mercaptoethanol, 1 mM TCEP). The fractions containing Sgs1 protein were concentrated on a 50 kDa MWCO Amicon concentrator and stored at −80°C in small aliquots.

#### Dmc1

Dmc1 was purified as described elsewhere (Busygina *et al*., 2013) with minor modifications. Briefly, the plasmid expressing Dmc1 protein with N-terminus (His)6-affinity tag (a kind gift from Lumir Krejci) was introduced into E. coli strain Rosetta(DE3)pLysS. Protein expression was induced by 0.5 mM IPTG at 37°C for 3 hours in LB media supplemented with ampicillin (100 μg/mL). The cell pellet was resuspended in the lysis buffer (25 mM Tris-HCl pH 7.5, 500 mM NaCl, 10% glycerol, 0.5 mM EDTA, 0.01% NP40, 1 mM DTT, 1 mM MgCl2, 1 mM ATP, and AEBSF). Resuspended cells were lysed by sonication before clearance at 35,000 rpm at 4°C for 45 min. Cleared lysate was incubated with 400 μl of Talon Resin (TaKaRa) for 1 hour at 4°C. The beads were washed with 10 mL of buffer T (25 mM Tris-HCl pH 7.5, 10% glycerol, 0.5 mM EDTA, 0.01% NP40, 1 mM DTT) containing 150 mM NaCl followed by additional washing step with 10 mL of buffer T containing 500 mM NaCl. The protein was eluted in steps with 200 and 500 mM imidazole in buffer T containing 140 mM NaCl, 1 mM MgCl2, and 1 mM ATP. Fractions containing Dmc1 protein were applied onto a 5-mL HiTrap Heparin HP affinity column (Cytiva) equilibrated with buffer T containing 140 mM NaCl, and eluted using a 25-mL gradient of 140-1000 mM NaCl in buffer T containing 1 mM MgCl2 and 1 mM ATP. The peak fractions were concentrated on a 30 kDa MWCO Amicon concentrator and applied onto a Superose 6 10/300 column (Cytiva) pre-equilibrated in SEC buffer (20 mM HEPES pH 7.5, 300 mM NaCl, 5% glycerol, 1 mM DTT) supplied with 1 mM MgCl2 and 1 mM ATP. The fractions containing Dmc1 were concentrated on a 30 kDa MWCO Amicon concentrator and stored at −80°C in small aliquots.

#### RPA

RPA complex was produced in E. coli strain C41 by coexpression of pCOLI-Twin-StrepII-Rfa1, pCDF-6xHis-Rfa2, and pRSF-6xHis-Rfa3 plasmids. Protein expression was induced by 0.5 mM IPTG at 25°C for 3 hours in TB media supplemented with ampicillin (100 μg/mL), kanamycin (25 μg/mL), and spectinomycin (50 μg/mL). The cell pellet was resuspended in the lysis buffer (50 mM HEPES pH 7.5, 300 mM NaCl, 5% glycerol, 0.01% NP40, 5 mM β-mercaptoethanol, AEBSF). Resuspended cells were lysed by sonication before clearance at 35,000 rpm at 4°C for 40 min. Cleared lysate was loaded on a 5 mL Strep-Tactin XT column (IBA) followed by wash using 25 mL of H buffer (20 mM HEPES pH 7.5, 5% glycerol, 0.01% NP40, 1 mM β-mercaptoethanol) containing 150 mM NaCl. The protein was eluted with a 50 mL of H buffer containing 100 mM NaCl and 50 mM biotin. Partially purified protein was further loaded onto a 5 mL HiTrap Heparin HP affinity column (Cytiva) pre-equilibrated in H buffer containing 150 mM NaCl and eluted with increasing salt gradient to 1 M NaCl. The peak fractions were concentrated on a 10 kDa MWCO Amicon concentrator and applied onto a Superose 6 10/300 column (Cytiva) pre-equilibrated in SEC buffer (20 mM HEPES pH 7.5, 300 mM NaCl, 5% glycerol, 1 mM β-mercaptoethanol, 1 mM TCEP). The fractions containing the RPA complex were concentrated on a 10 kDa MWCO Amicon concentrator and stored at −80°C in small aliquots.

### Mass Photometry

Mass Photometry was performed in the mass photometry buffer (MP) containing 30 mM HEPES pH 7.8, 150 mM NaCl, 5% glycerol, 1 mM MgCl2, and 1 mM TCEP. Protein samples (3 μM) were pre-equilibrated for 1 hour in the MP buffer. Measurements were performed using Refeyn One (Refyn Ltd., Oxford, UK) mass photometer. Directly before the measurement, the sample was diluted 1:100 with the MP buffer. Molecular mass was determined in Analysis software provided by the manufacturer using a NativeMark (Invitrogen) based standard curve created under the identical buffer composition.

### DNA substrates

Fluorescently labelled DNA substrates were prepared as described previously (De Muyt *et al*., 2018; Ranjha *et al*., 2014). The individual DNA substrates were prepared by annealing 5^*t*^ fluorescently labelled oligonucleotide 1253 (or 1253-T) with the following oligonucleotides: dsDNA (1253, 1253C); 3^*t*^overhang (1253, 3^*t*^overhang25nt); Y-form (1253-T, 1254); D-loop (1253-T, 315, 320, X12-3SC), HJ (1253-T, 1254, 1255, 1256). The sequences of all oligonucleotides used in this study are listed in Supplementary Table 2.

### Electrophoretic mobility shift assays (EMSAs)

The binding reactions (10 μL volume) were carried out in EMSA buffer (25 mM HEPES pH 7.5, 0.1 μg/μL BSA, 60 mM NaCl) containing indicated fluorescently labelled DNA substrate (10 nM). The reactions were started by addition of increasing amounts of Mer3 protein (37.5, 75, 150, and 300 nM) and incubated for 20 min at 30 °C. After the addition of 2 μL of the gel loading buffer (60% glycerol, 10 mM Tris-HCl, pH 7.4, 60 mM EDTA, 0.15 % Orange G), the reaction mixtures were resolved in 0.8% agarose gel in 1x TAE buffer (40 mM Tris, 20 mM acetic acid, 1 mM EDTA). The gels were scanned using Amersham Typhoon scanner (Cytiva) and quantified by ImageQuant TL software (Cytiva).

### Strand separation assays

The strand separation assays (10 μL volume) were carried out in SS buffer (25 mM HEPES pH 7.5, 60 mM NaCl, 0.1 μg/μL BSA, 1 mM MgCl2, 1 mM ATP, 10 mM creatine phosphate, 15 μg/ml creatine kinase) containing indicated fluorescently labelled DNA substrate (5 nM). The reactions were started by addition of increasing amounts of Mer3 protein (10, 20, 40, and 80 nM). After the incubation for 30 min at 30 °C the reactions were stopped with 0.5 mg/mL proteinase K and 0.1% SDS, and incubated for 5 min at 37°C. The samples were then mixed with 2 μL of the gel loading buffer (60% glycerol, 10 mM Tris-HCl, pH 7.4, 60 mM EDTA) and separated on 10% (w/v) native polyacrylamide gel in 1xTBE buffer at a constant voltage of 110 V for 1 hour at 4°C. The gels were scanned using Amersham Typhoon scanner (Cytiva) and quantified by ImageQuant TL software (Cytiva).

### Cross-linking Mass Spectrometry (XL-MS)

For XL-MS analysis proteins were diluted in 200 μl of XL-MS buffer (30 mM HEPES 6.8, 150 mM NaCl, 5% glycerol, 1 mM MgCl2, 1 mM TCEP) to the final concentration of 3 μM, mixed with 3 μL of DSBU (200 mM) and incubated for 1 hour at 25°C. The reaction was stopped by adding 20 μL of 1 M Tris-HCl pH 8.0 and incubated for another 30 min at 25°C. The crosslinked sample was precipitated by addition of 4X volumes of 100% cold acetone followed by overnight incubation at −20°C. Samples were analysed as previously described (Pan *et al*., 2018). For interaction network visualisation XVis software was used and for visualisation of the crosslinks on the PDB model PyXlinkViewer (Schiffrin *et al*., 2020) and XMAS (Lagerwaard *et al*., 2022) was used. Each time a different cutoff for the cross-linking credibility was selected depending on the quality of the cross-linking data.

### SEC-MALS

50 μL samples at 10 μM concentration were loaded onto a Superose 6 5/150 (for fhe full length protein) analytical size exclusion column (Cytiva) equilibrated in SEC-MALS buffer (30 mM HEPES pH 6.8, 300 mM NaCl, 1 mM MgCl2, 5 μM ZnCl2, 1 mM TCEP) attached to a 1260 Infinity II LC System (Agilent). MALS was carried out using a Wyatt DAWN detector attached in line with the size exclusion column. Mer3 fragments were analysed on Superdex 200 5/150 column (Cytiva) equilibrated in SEC-MALS2 buffer (50 mM HEPES pH 7.5, 300 mM NaCl, 1 mM TCEP).

### Alphafold2 Predictions

The predicted structure of full-length Mer3 was obtained from the publicly available database (https://www.alphafold.ebi.ac.uk/ ID P51979). The predicted structures of Mlh1/Mlh2, Top3/Rmi1 and Sgs1/Top3/Rmi1 were calculated using AlphaFold Multimer (2.2.0) (Evans *et al*., 2021) run on GPU nodes of the Raven HPC of the Max Planck Computing and Data Facility (MPCDF), Garching. Each job was run on a single node consisting of 4 x Nvidia A100 NVlink 40 GB GPUs. Multiple predictions were generated for each run, and the best model (determined by pTM score) was then used. PAE plots were generated using a custom script (Vikram Alva, MPI Biology Tübingen).

### Pull-down assays

For pull-down between Mer3 and Top3/Rmi1, GST-tagged Top3/Rmi1 (3 μg) was incubated with Strep-tagged Mer3 (3 μg) in the reaction buffer (25 mM HEPES pH 7.5, 5% glycerol, 100 mM NaCl, 1 mM TCEP, 0.1% Tween-20) for 20 min at 30°C in the thermomixer (950 rpm). Pre-washed magnetic glutathione beads (1 μL) were then added to the samples and the mixtures were incubated for 2 min at 30°C in the thermomixer (750 rpm). Beads were washed twice with 100 μL of the reaction buffer. The proteins were eluted by boiling in 30 μL 2x SDS Laemmli buffer. The samples were loaded onto 10% SDS-PAGE and stained by Der Blaue Jonas gel dye.

For pull-down between Top3 and Mer3, His-tagged Top3 (3 μg) was incubated with Strep-tagged Mer3 (3 μg) in the reaction buffer (25 mM HEPES pH 7.5, 5% glycerol, 50 mM NaCl, 1 mM TCEP, 0.1% Tween-20) for 20 min at 30°C in the thermomixer (950 rpm). Anti-Strep antibody (2 μL, Abcam, ab76949) was added to the reactions and the mixtures were incubated for 1 hour at 4°C in the thermomixer (950 rpm). Finally, 0.5 μL pre-washed magnetic-conjugated protein G beads (Dynabeads protein G, Invitrogen) were added to the reactions followed by the incubation for 1 hour at 4°C in the thermomixer (950 rpm). Beads were washed twice with 100 μL of the reaction buffer. The proteins were eluted by boiling in 30 μL 2x SDS Laemmli buffer. The samples were loaded onto 11% SDS-PAGE and analysed by western blot.

For pull-down between Rmi1 and Mer3, GST-tagged Rmi1 (4 μg) was incubated with Strep-tagged Mer3 (4 μg) in the reaction buffer (25 mM HEPES pH 7.5, 5% glycerol, 75 mM NaCl, 1 mM TCEP, 0.1% Tween-20) for 20 min at 30°C in the thermomixer (950 rpm). Pre-washed glutathione beads (10 μL) were then added to the samples and the mixtures were incubated for 30 min at 7°C in the thermomixer (950 rpm). Beads were washed twice with 100 μL of the reaction buffer. The proteins were eluted by boiling in 30 μL 2x SDS Laemmli buffer. The samples were loaded onto 11% SDS-PAGE and stained with coomassie brilliant blue.

For pull-down between Mer3 and Top3/Rmi1 in the presence or absence of Sgs1(1-107), GST-tagged Top3/Rmi1 (4 μg) was incubated with Strep-tagged Mer3 (4 μg) in the reaction buffer (25 mM HEPES pH 7.5, 5% glycerol, 100 mM NaCl, 1 mM TCEP, 0.1% Tween-20) for 20 min at 30°C in the thermomixer (950 rpm). For competition assays, increasing amounts of His-tagged Sgs1(1-107) were added to the reactions. Pre-washed glutathione beads (10 μL) were then added to the samples and the mixtures were incubated for 30 min at 7°C in the thermomixer (950 rpm). Beads were washed twice with 100 μL of the reaction buffer. The proteins were eluted by boiling in 30 μL 2x SDS Laemmli buffer. The samples were loaded onto 11% or 13% SDS-PAGE and stained with coomassie brilliant blue.

For pull-down between de-/phosphorylated variants of Mer3 and Top3/Rmi1 in the presence or absence of Sgs1(1-605), GST-tagged Top3/Rmi1 (3 μg) was incubated with Strep-tagged Mer3 (3 μg) in the reaction buffer (25 mM HEPES pH 7.5, 5% glycerol, 100 mM NaCl, 1 mM TCEP, 0.1% Tween-20) for 20 min at 30°C in the thermomixer (950 rpm). For competition assays, increasing amounts of untagged Sgs1(1-605) were added to the reactions. Pre-washed magnetic glutathione beads (1 μL) were then added to the samples and the mixtures were incubated for 2 min at 30°C in the thermomixer (750 rpm). Beads were washed twice with 100 μL of the reaction buffer. The proteins were eluted by boiling in 30 μL 2x SDS Laemmli buffer. The samples were loaded onto 10% SDS-PAGE and stained by Der Blaue Jonas gel dye.

For pull-down between Mlh1-Mlh2 and Mer3, MBP-tagged Mlh1-Mlh2 complex (1 μg) was incubated with Strep-tagged Mer3 (1 μg) in the reaction buffer (25 mM HEPES pH 7.5, 100 mM NaCl, 1 mM MgCl2, 1 mM DTT) for 20 min at 30°C in the thermomixer (950 rpm). Anti-MBP antibody (0.5 μL, Invitrogen, PA1-989) was added to the reactions and the mixtures were incubated for 1 hour at 4°C in the thermomixer (750 rpm). Finally, 1 μL pre-washed magnetic-conjugated protein G beads (Dynabeads protein G, Invitrogen) were added to the reactions followed by the incubation for 1 hour at 4°C in the thermomixer (750 rpm). Beads were washed twice with 100 μL of the reaction buffer. The proteins were eluted by boiling in 30 μL 2x SDS Laemmli buffer. The samples were loaded onto 9% SDS-PAGE and analysed by western blot. Pull-down between Mer3, Mlh1-Mlh2 and Top3/GST-Rmi1 complex (all 1 μg) was done using the same protocol but in reaction buffer containing 25 mM HEPES pH 7.5, 5% glycerol, 75 mM NaCl, 1 mM TCEP, 0.1% Tween-20.

### Microscale thermophoresis (MST)

Binding affinity analysis by microscale thermophoresis was performed using the Monolith NT instrument (Nanotemper Technologies). All reactions (in triplicates) were done in the commercial MST buffer (50 mM Tris-HCl pH 7.4, 150 mM NaCl, 10 mM MgCl2; Nanotemper Technologies) supplied with 0.05% Tween-20. Measurements were performed at 25°C, and contained constant concentration of 45 nM RED-NHS labelled Mer3 (labelling was performed according to the manufacturer’s protocol - Nanotemper Technologies) and increasing concentrations of Top3/Rmi1, Rmi1, or Mlh1/Mlh2, respectively. Data were analysed by the MO.Affinity Analysis software (NanoTemper Technologies).

### D-loop assay

The reactions (in a total volume 22 μL) were performed in D-loop reaction buffer (25 mM HEPES pH 7.5, 0.1 μg/μL BSA, 100 mM NaCl, 1 mM MgCl2, 1 mM ATP, 10 mM creatine phosphate, 15 μg/ml creatine kinase). Radioactively labelled 90-mer ssDNA (oWL981, 2 μM nucleotides) was incubated with Dmc1 protein (1.5 μM) for 5 min at 37°C followed by addition of RPA (90 nM) and additional incubation for 5 min at 37°C. The formation of D-loop was started by addition of pUC19 plasmid (18 nM molecules). After 15 min incubation at 30°C, Sgs1 (15 nM), Top3/Rmi1 (15 nM) and the increasing amounts of Mer3 and/or Mlh1/Mlh2 (15, 150, 300 nM) were added to the reactions. At the indicated time points, 10.5 μL of sample was mixed with 0.5% SDS (final) and 0.5 mg/mL proteinase K followed by incubation for 15 min at 37°C. The deproteinized samples were separated in a 0.9% agarose gel. After electrophoresis, the gel was dried on grade 3 chromosome paper (Whatman), exposed to a phosphorimager screen, and visualised using Amersham Typhoon scanner (Cytiva). The quantification was done using ImageQuant TL software (Cytiva).

### Yeast strains

All strains, except those used for Y2H analysis, were derived from Saccharomyces cerevisiae SK1 strains YML4068 and YML4069 (a kind gift from Joao Matos) and their genotypes are listed in Supplementary Table 3. C-terminal tagging of Mer3 and Top3 was done using the PCR-based method as previously described (Janke *et al*., 2004; Knop *et al*., 1999). Mer3 was tagged with 9 copies of Myc tag by using plasmid pYM18. Top3 was tagged by 3 copies of HA tag using the plasmid pYM24. The correct epitope tag insertion was confirmed by PCR.

### Yeast two-hybrid

Yeast genes ORFs were PCR-amplified from SK1 strain genomic DNA. MER3 was prepared by Gibson assembly of 2 PCR products eliminating MER3’s intron. The corresponding genes were cloned into pGAD-C1 or pGBDU-C1 vectors, respectively. The resulting plasmids were co-transformed into the *S. cerevisiae* reporter strain (yWL365; a kind gift from Gerben Vader) and plated onto the selective medium lacking leucine and uracil. For drop assay, 2.5 μL from 10-fold serial dilutions of cell cultures with the initial optical density (OD600) of 0.5 were spotted onto -Leu/-Ura (control) and -Leu/-Ura/-His plates with or without 1 mM 3-aminotriazole. Cells were grown at 30°C for up to 4-6 days. and imaged.

### Meiotic time course

Cells were grown overnight in liquid YPD culture at 30°C followed by inoculation in pre-sporulation media (BYTA; 50 mM potassium phthalate, 1% yeast extract, 2% bacto tryptone, and 1% potassium acetate) at OD600 = 0.3 for additional 16-18 hours at 30°C. Next morning, cells were washed twice with sporulation medium (SPO, 0.3% potassium acetate) and resuspended in sporulation medium at OD600 = 1.9 to induce meiosis at 30°C.

### *In vivo* co-immunoprecipitation

100 mL of meiotic cultures (at 6 hours into a meiotic time course) were harvested by spinning down at 3,000 rpm for 5 min followed by washing with 500 μL of cold H2O containing 1 mM PMSF. Cell pellets were resuspended in 350 μL of ice-cold co-IP buffer (50 mM Tris-HCl pH 7.5, 150 mM NaCl, 1% Nonidet P-40, 1 mM EDTA pH 8.0, 1 mM PMSF, AEBSF, Serva protease cocktail and a cocktail of protease inhibitors which was freshly added) and glass beads. The cells were lysed using a FastPrep-24 disruptor (MP Biomedicals) (setting: 2x 40 sec cycles at speed 6.0). Lysates were cleared by 2 rounds of centrifugation for 10 min at 15,000 rpm and the supernatants were after each centrifugation step transferred to a clean microcentrifuge tube. 1 μL of antibody (anti-HA; Sigma-Aldrich H6908) was added to the samples followed by 3 hours incubation at 4°C. Subsequently, 25 μL of buffer-washed Dynabeads Protein G (Thermo Fisher Scientific) was added and the samples were incubated overnight at 4°C. The next day, Dynabeads were washed four times with 500 μL of ice-cold IP buffer. For the final wash, beads were transferred to a new microcentrifuge tube and washed 500 μL of ice-cold IP buffer without Nonidet P-40. The beads were resuspended in 55 μL of 2x SDS Laemmli buffer and incubated for 5 min at 95°C. The samples were loaded onto a 9% SDS-polyacrylamide gel and blotted to nitrocellulose membrane. Antibodies used were as follows: anti-PGK1 (22C5D8, Thermo Fisher Scientific, 459250, 1:1,000), anti-HA (Sigma-Aldrich, H6908, 1:1,000), anti-Myc (Abcam, ab1326, 1:1,000), goat anti-rabbit IgG peroxidase conjugate (Merck, 401353), goat anti-mouse IgG peroxidase conjugate (Merck, 401215). Signal was detected using ECL Prime Western Blotting Detection Reagents (Cytiva) and visualised by a ChemiDocMP (Bio-Rad Inc).

## Acknowledgements

We thank members of the Weir Lab for critical reading and input into the manuscript. IP-MS analysis was carried out in the Proteomics Facility at EMBL, Heidelberg. Thanks to Vikram Alva (MPI for Biology Tübingen) for advice on AlphaFold2 modelling and analysis. Thanks to Maria Kharlamova (Cellular Nanoscience, University of Tübingen) for her advice on the Mass Photometry experiments. We thank Gerben Vader (Cancer Centre Amsterdam), João Matos (Max Perutz Labs Vienna), and Lumír Krejčí (Masaryk University, Brno) for plasmids and yeast strains. Work in the Weir Lab is supported by the Max Planck Society and the German Research Foundation (Grant Number WE 6513/2-1).

## CRediT author statement

Veronika Altmannova: Methodology, Validation, Formal analysis, Investigation, Writing - Original Draft, Visualisation. Magdalena Firlej: Methodology, Formal analysis, Investigation, Writing - Original Draft, Visualisation. Franziska Mller: Methodology, Investigation, Petra Janning: Methodology, Formal analysis. Rahel Rauleder: Formal analysis, Investigation. Dorota Rousova: Formal analysis, Investigation. Andreas Schler: Investigation. John R. Weir: Conceptualization, Formal analysis, Writing - Original Draft, Writing - Review Editing, Visualisation, Supervision, Project administration, Funding acquisition

## Supplementary Data

**Supplementary table 1.** Plasmids used in this study.

| Plasmid ID | Description | Reference |
| --- | --- | --- |
| pWL413 | pLIB-Mer3-Strep | This study |
| pWL522 | pCOLI-Strep-Rfa1 | This study |
| pWL526 | pCDF-6xHis-Rfa2 | This study |
| pWL527 | pRSF-6xHis-Rfa3 | This study |
| pWL746 | pET11c-Dmc1 | Lumir Krejci |
| pWL808 | pLIB-GST-Rmi1 | This study |
| pWL812 | pLIB-6xHis-Top3 | This study |
| pWL856 | pBIG1a-Top3/GST-Rmi1 | This study |
| pWL993 | pBIG1a-6xHis-MBP-Mlh1 6xHis-MBP-Mlh2 | This study |
| pWL1645 | pBIG1a-6xHis-Mlh1 Strep-Mlh2 | This study |
| pWL1016 | pLIB-6xHis-MBP-Sgs1-6xHis | This study |
| pWL1097 | pLIB-Mer3(K167A)-Strep | This study |
| pWL1758 | pCOLI-6xHis-Sgs1(1-107) | This study |
| pWL1897 | pCOLI-6xHis-MBP-Sgs1(1-605) | This study |
| pWL1565 | pGAD-C1 | This study |
| pWL1564 | pGBDU-C1 | This study |
| pWL1700 | pGAD-C1-Mer3 | This study |
| pWL1716 | pGBDU-C1-Mer3 | This study |
| pWL1713 | pGAD-C1-Mer3(1-1023) | This study |
| pWL1712 | pGBDU-C1-Mer3(1-1023) | This study |
| pWL1826 | pGAD-C1-Mer3(122-1187) | This study |
| pWL1827 | pGBDU-C1-Mer3(122-1187) | This study |
| pWL1696 | pGBDU-C1-Top3 | This study |
| pWL1695 | pGBDU-C1-Rmi1 | This study |

**Supplementary Table 2.**
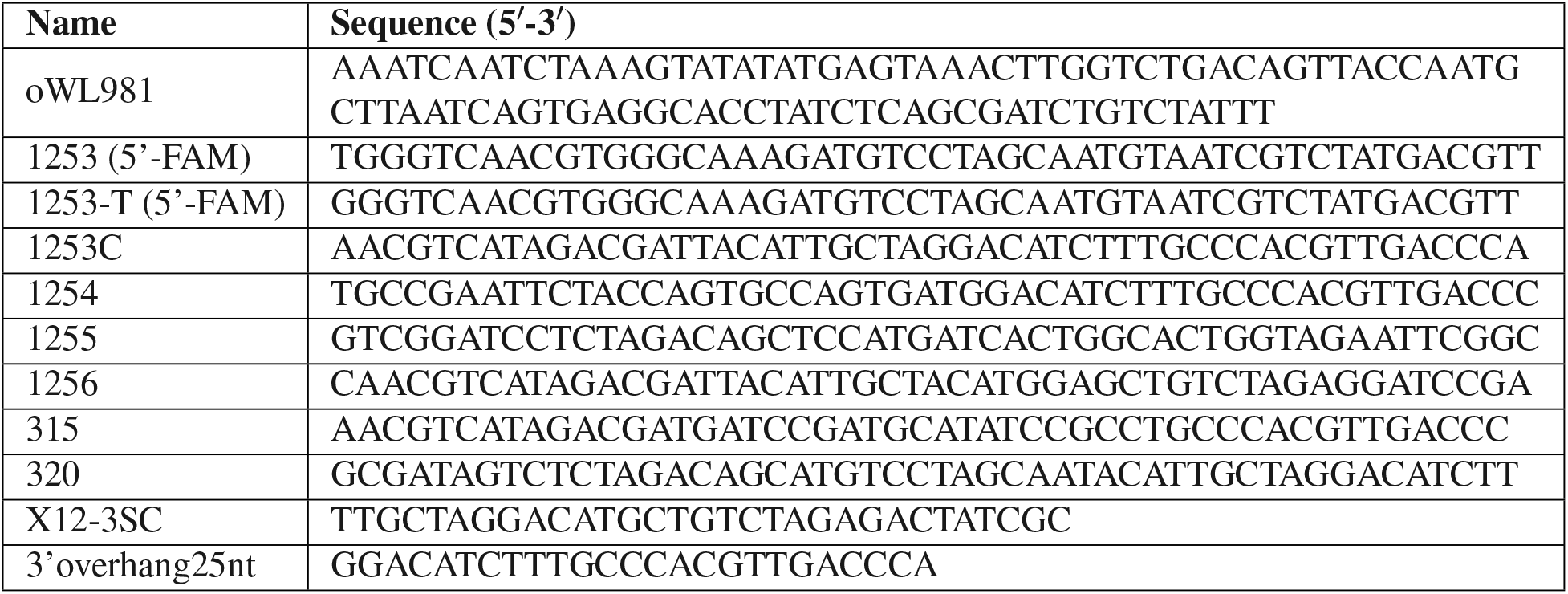
Oligonucleotides used in this study.

**Supplementary Table 3.**
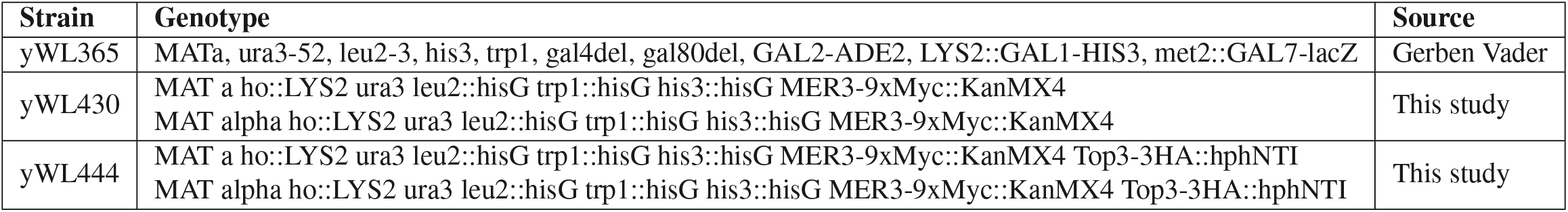
Yeast strains used in this study.

| Strain | Genotype | Source |
| --- | --- | --- |
| yWL365 | MATa, ura3-52, leu2-3, his3, trp1, gal4del, gal80del, GAL2-ADE2, LYS2::GAL1-HIS3, met2::GAL7-lacZ | Gerben Vader |
| yWL430 | MAT a ho::LYS2 ura3 leu2::hisG trp1::hisG his3::hisG MER3-9xMyc::KanMX4<br>MAT alpha ho::LYS2 ura3 leu2::hisG trp1::hisG his3::hisG MER3-9xMyc::KanMX4 | This study |
| yWL444 | MAT a ho::LYS2 ura3 leu2::hisG trp1::hisG his3::hisG MER3-9xMyc::KanMX4 Top3-3HA::hphNTI<br>MAT alpha ho::LYS2 ura3 leu2::hisG trp1::hisG his3::hisG MER3-9xMyc::KanMX4 Top3-3HA::hphNTI | This study |

**Supplementary Figure 1.**
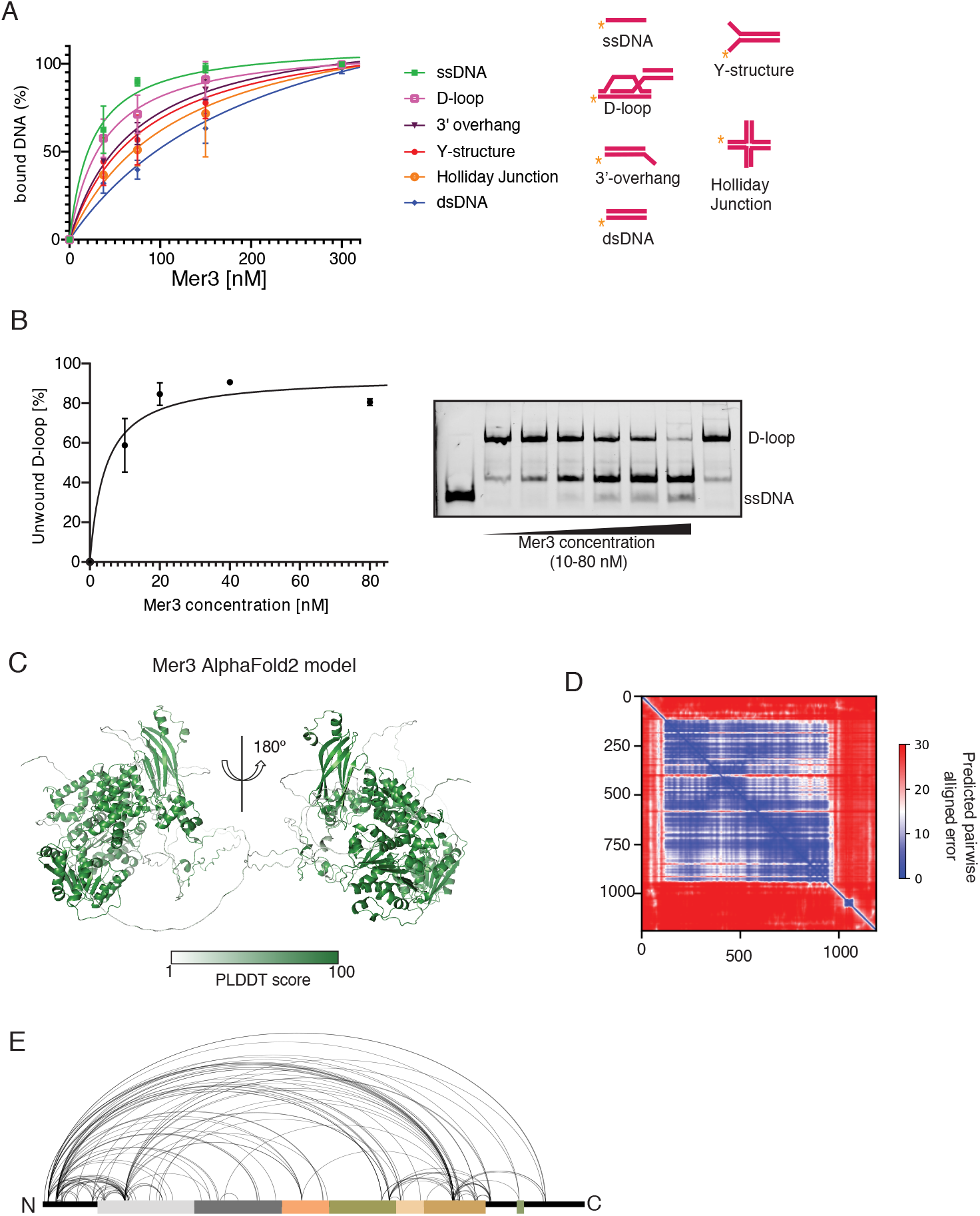
A) Summary of EMSAs of recombinant Mer3-Strep binding to different substrates (cartoon representations of substrates, right). Mer3 bound to ssDNA and D-loops with the highest affinity. B) Strand separation activity of Mer3 on D-loop substrate. Graph shows quantification of three independent experiments (example shown right) for strand separation activity. C) AlphaFold2 model of S. cerevisiae Mer3 coloured according to pLDDT score. D) PAE plot of the AlphaFold2 model of Mer3. E) Visualisation of Mer3 XL-MS data showing the intrachain cross-links on the domain cartoon of Mer3 (domain cartoon coloured as in Figure 1D

**Supplementary Figure 2.**
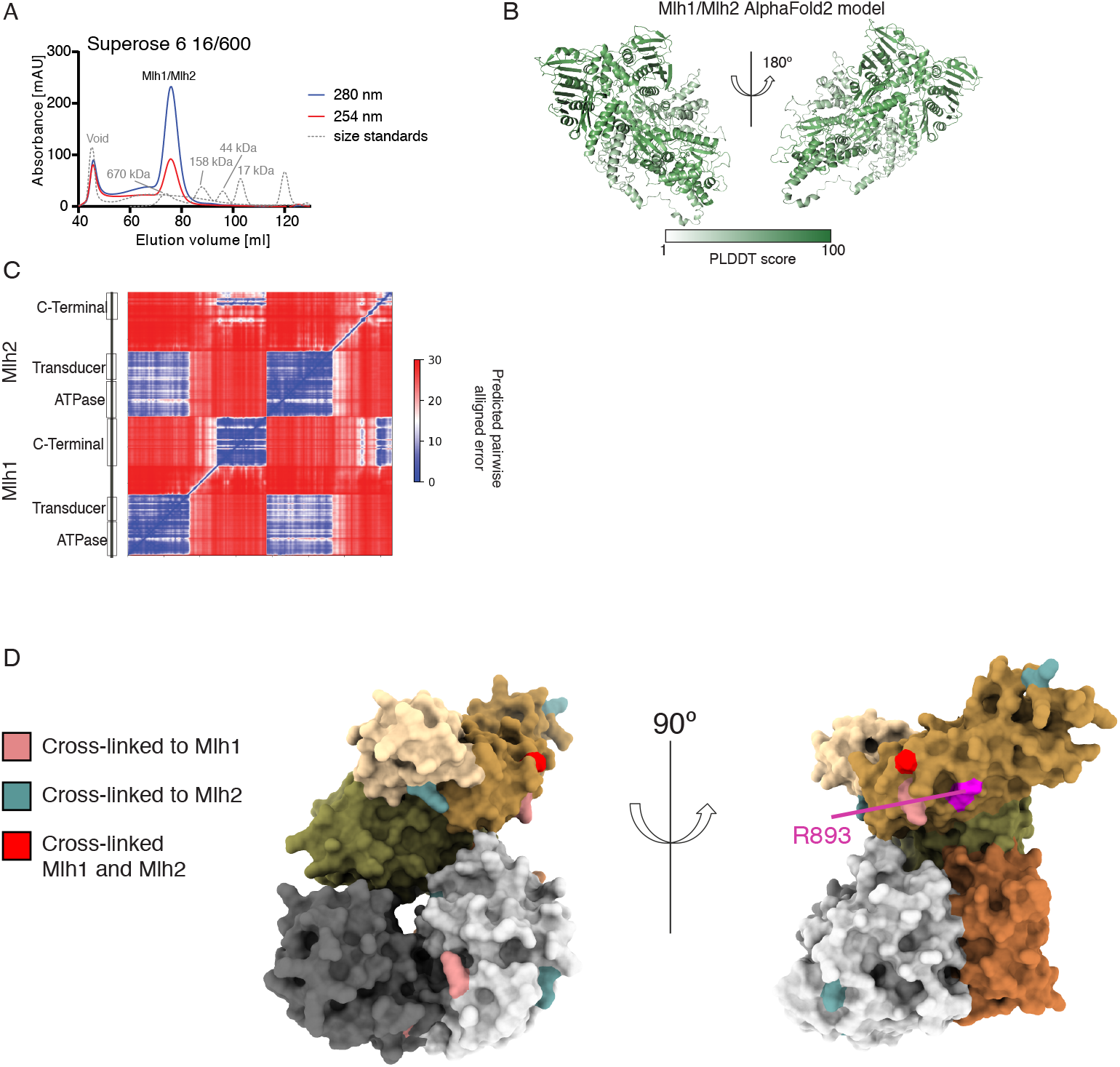
A) SEC profile of the purification of S. cerevisiae Mlh1/Mlh2 showing absorbance at 280 nm (blue) or 260 nm (red). SEC standards include the bio-rad gel filtration standard, and dextran blue as a void marker. B) AF2 multimer structure of Mlh1/Mlh2 heterodimer coloured by the pLDDT score. C) PAE plot of the AF2 multimer prediction of the Mlh1/Mlh2 heterodimer D) Surface representation of the AF2 model of Mer3. Domains coloured as in Figure 1D. Residues coloured additionally according to which cross-links detected within the context of the DSBU treated Mer3/Mlh1/Mlh2 complex (Mlh1, pink; Mlh2, blue; both, red). Location of R893 is highlighted as this was previously shown to disrupt the interaction between Mer3 and Mlh1/Mlh2.

**Supplementary Figure 3.**
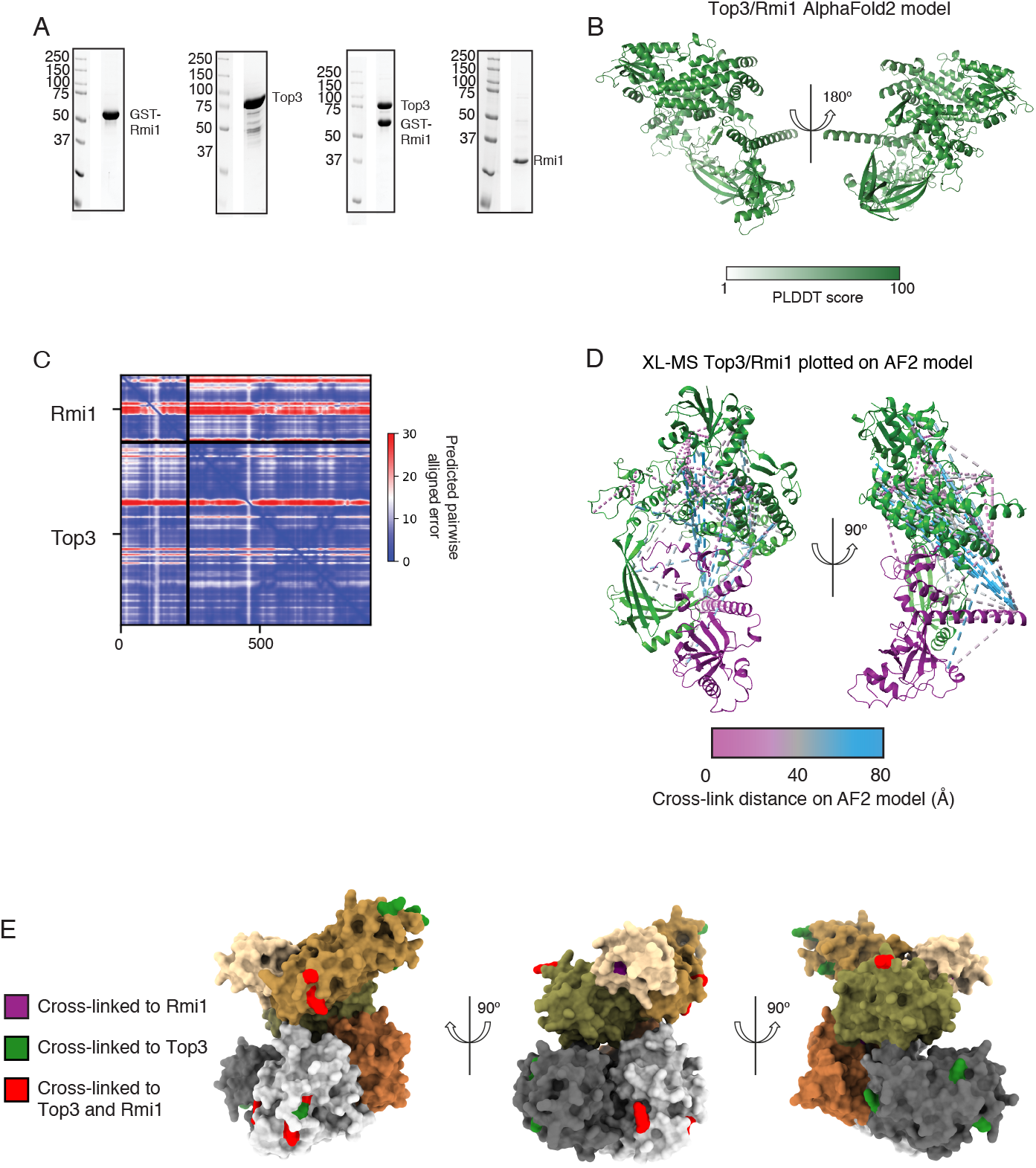
A) Representative coomassie stained SDS-PAGE lanes from purifications of GST-Rmi1, His-Top3, Top3/GST-Rmi1 and Top3/Rmi1. B) AF2 multimer structure of *S. cerevisiae* Top3/Rmi1 heterodimer coloured by the pLDDT score. C) PAE plot of the AF2 multimer structure of S. cerevisiae Top3/Rmi1 heterodimer. D) XL-MS data of the Top3/Rmi1 complex plotted onto the AlphaFold2 multimer model using XMAS (Lagerwaard et al., 2022). Cross-links are coloured according to distance. E) Surface representation of the AF2 model of Mer3. Domains coloured as in Figure 1D. Residues coloured additionally according to which cross-links detected within the context of the DSBU treated Mer3/Top3/Rmi1 complex (Top3, green; Rmi1, purple; both, red).

**Supplementary Figure 4.**
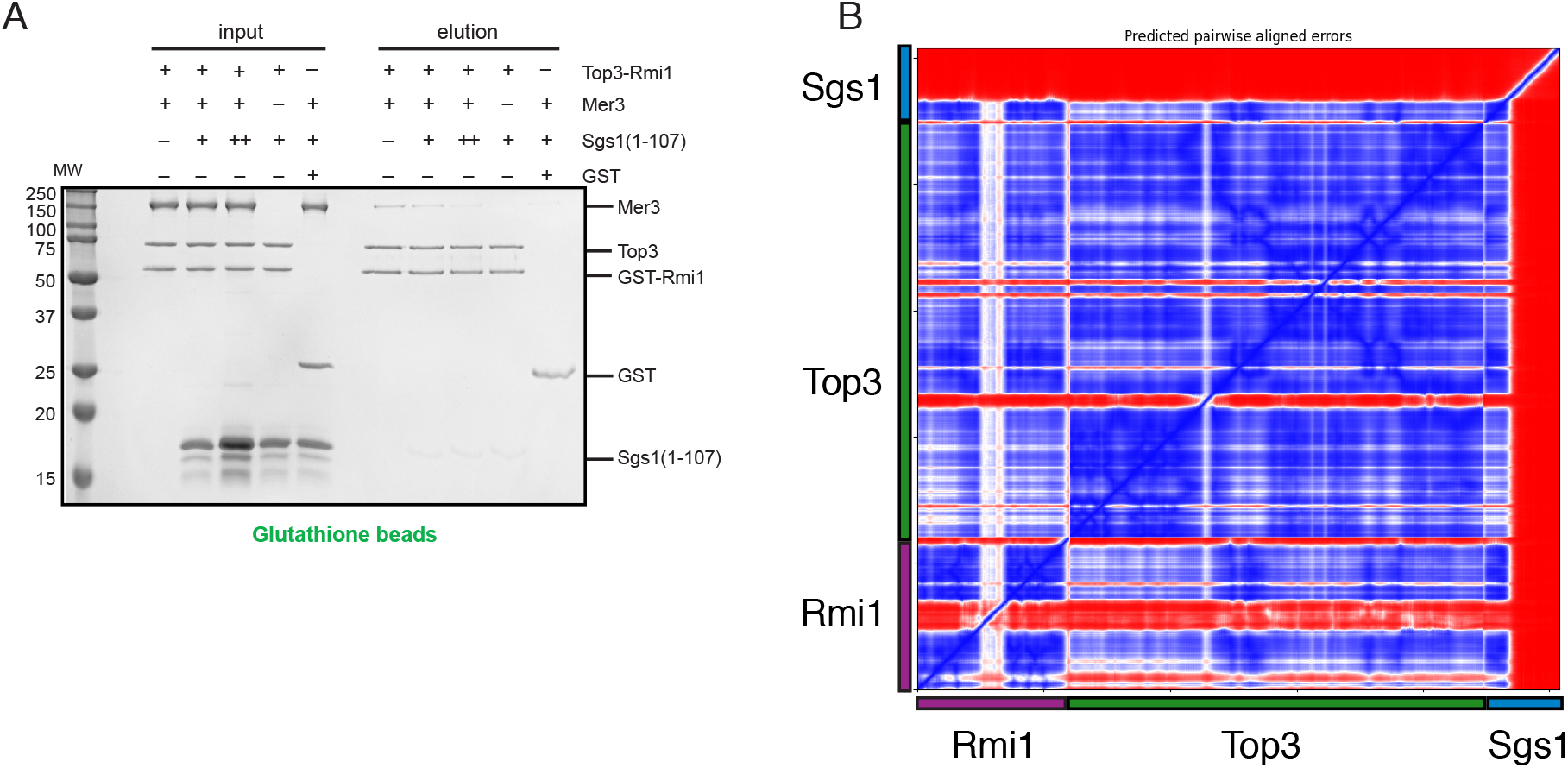
A) Glutathione pulldown of recombinant Top3/GST-Rmi1 against Mer3-Strep, and in increasing concentrations of Sgs11-107. GST alone is used as a control for background binding. B) PAE plot of the AF2 multimer predicted structure of Sgs11-107/Top3/Rmi1.

## References

Altmannova, V., Blaha, A., Astrinidis, S., Reichle, H., and Weir, J.R. (2021). InteBac: An integrated bacterial and baculovirus expression vector suite. Protein Sci. 30, 108–114. https://doi.org/10.1002/pro.3957.

Amin, A.D., Chaix, A.B.H., Mason, R.P., Badge, R.M., and Borts, R.H. (2010). The roles of the Saccharomyces cerevisiae RecQ helicase SGS1 in meiotic genome surveillance. PLoS One 5, e15380. https://doi.org/10.1371/journal.pone.0015380.

Bachrati, C.Z., Borts, R.H., and Hickson, I.D. (2006). Mobile D-loops are a preferred substrate for the Bloom’s syndrome helicase. Nucleic Acids Res. 34, 2269–2279. https://doi.org/10.1093/nar/gkl258.

Ban, C., Junop, M., and Yang, W. (1999). Transformation of MutL by ATP binding and hydrolysis: a switch in DNA mismatch repair. Cell 97, 85–97. https://doi.org/10.1016/s0092-8674(00)80717-5.

Bennett, R.J., Noirot-Gros, M.-F., and Wang, J.C. (2000). Interaction between Yeast Sgs1 Helicase and DNA Topoisomerase III *. J. Biol. Chem. 275, 26898–26905. https://doi.org/10.1016/S0021-9258(19)61459-6.

Bernstein, K.A., Gangloff, S., and Rothstein, R. (2010). The RecQ DNA helicases in DNA repair. Annu. Rev. Genet. 44, 393–417. https://doi.org/10.1146/annurev-genet-102209-163602.

Bieniossek, C., Imasaki, T., Takagi, Y., and Berger, I. (2012). MultiBac: expanding the research toolbox for multiprotein complexes. Trends Biochem. Sci. 37, 49–57. https://doi.org/10.1016/j.tibs.2011.10.005.

Bizard, A.H., and Hickson, I.D. (2014). The dissolution of double Holliday junctions. Cold Spring Harb. Perspect. Biol. 6, a016477. https://doi.org/10.1101/cshperspect.a016477.

Bocquet, N., Bizard, A.H., Abdulrahman, W., Larsen, N.B., Faty, M., Cavadini, S., Bunker, R.D., Kowalczykowski, S.C., Cejka, P., Hickson, I.D., et al. (2014). Structural and mechanistic insight into Holliday-junction dissolution by topoisomerase III? and RMI1. Nat. Struct. Mol. Biol. 21, 261–268. https://doi.org/10.1038/nsmb.2775.

Börner, G.V., Kleckner, N., and Hunter, N. (2004). Crossover/noncrossover differentiation, synaptonemal complex formation, and regulatory surveillance at the leptotene/zygotene transition of meiosis. Cell 117, 29–45. https://doi.org/10.1016/s0092-8674(04)00292-2.

van Brabant, A.J., Ye, T., Sanz, M., German, J.L., III, Ellis, N.A., and Holloman, W.K. (2000). Binding and melting of D-loops by the Bloom syndrome helicase. Biochemistry 39, 14617–14625. https://doi.org/10.1021/bi0018640.

Cejka, P., Plank, J.L., Bachrati, C.Z., Hickson, I.D., and Kowalczykowski, S.C. (2010). Rmi1 stimulates decatenation of double Holliday junctions during dissolution by Sgs1-Top3. Nat. Struct. Mol. Biol. 17, 1377–1382. https://doi.org/10.1038/nsmb.1919.

Cejka, P., Plank, J.L., Dombrowski, C.C., and Kowalczykowski, S.C. (2012). Decatenation of DNA by the S. cerevisiae Sgs1-Top3-Rmi1 and RPA complex: a mechanism for disentangling chromosomes. Mol. Cell 47, 886–896. https://doi.org/10.1016/j.molcel.2012.06.032.

Chang, M., Bellaoui, M., Zhang, C., Desai, R., Morozov, P., Delgado-Cruzata, L., Rothstein, R., Freyer, G.A., Boone, C., and Brown, G.W. (2005). RMI1/NCE4, a suppressor of genome instability, encodes a member of the RecQ helicase/Topo III complex. EMBO J. 24, 2024–2033. https://doi.org/10.1038/sj.emboj.7600684.

Chen, C., Zhang, W., Timofejeva, L., Gerardin, Y., and Ma, H. (2005). The Arabidopsis ROCK-N-ROLLERS gene encodes a homolog of the yeast ATP-dependent DNA helicase MER3 and is required for normal meiotic crossover formation. Plant J. 43, 321–334. https://doi.org/10.1111/j.1365-313X.2005.02461.x.

De Muyt, A., Jessop, L., Kolar, E., Sourirajan, A., Chen, J., Dayani, Y., and Lichten, M. (2012). BLM helicase ortholog Sgs1 is a central regulator of meiotic recombination intermediate metabolism. Mol. Cell 46, 43–53. https://doi.org/10.1016/j.molcel.2012.02.020.

De Muyt, A., Pyatnitskaya, A., Andréani, J., Ranjha, L., Ramus, C., Laureau, R., Fernandez-Vega, A., Holoch, D., Girard, E., Govin, J., et al. (2018). A meiotic XPF-ERCC1-like complex recognizes joint molecule recombination intermediates to promote crossover formation. Genes Dev. 32, 283–296. https://doi.org/10.1101/gad.308510.117.

Duroc, Y., Kumar, R., Ranjha, L., Adam, C., Guérois, R., Md Muntaz, K., Marsolier-Kergoat, M.-C., Dingli, F., Laureau, R., Loew, D., et al. (2017). Concerted action of the MutL? heterodimer and Mer3 helicase regulates the global extent of meiotic gene conversion. Elife 6, e21900. https://doi.org/10.7554/eLife.21900.

Evans, R., O’Neill, M., Pritzel, A., Antropova, N., Senior, A., Green, T., Žk, A., Bates, R., Blackwell, S., Yim, J., et al. (2021). Protein complex prediction with AlphaFold-Multimer.

Grigaitis, R., Ranjha, L., Wild, P., Kasaciunaite, K., Ceppi, I., Kissling, V., Henggeler, A., Susperregui, A., Peter, M., Seidel, R., et al. (2020). Phosphorylation of the RecQ Helicase Sgs1/BLM Controls Its DNA Unwinding Activity during Meiosis and Mitosis. Dev. Cell 53, 706-723.e5. https://doi.org/10.1016/j.devcel.2020.05.016.

Grimm, M., Zimniak, T., Kahraman, A., and Herzog, F. (2015). xVis: a web server for the schematic visualization and interpretation of crosslink-derived spatial restraints. Nucleic Acids Research 43, W362–W369. https://doi.org/10.1093/nar/gkv463.

Guiraldelli, M.F., Eyster, C., Wilkerson, J.L., Dresser, M.E., and Pezza, R.J. (2013). Mouse HFM1/Mer3 is required for crossover formation and complete synapsis of homologous chromosomes during meiosis. PLoS Genet. 9, e1003383. https://doi.org/10.1371/journal.pgen.1003383.

Holm, L. (2020). Using Dali for Protein Structure Comparison. Methods Mol. Biol. 2112, 29–42. https://doi.org/10.1007/978-1-0716-0270-6.

Humphryes, N., and Hochwagen, A. (2014). A non-sister act: Recombination template choice during meiosis. Exp. Cell Res. 329, 53–60. https://doi.org/10.1016/j.yexcr.2014.08.024.

Hunter, N., and Kleckner, N. (2001). The single-end invasion: an asymmetric intermediate at the double-strand break to double-holliday junction transition of meiotic recombination. Cell 106, 59–70. https://doi.org/10.1016/s0092-8674(01)00430-5.

Janke, C., Magiera, M.M., Rathfelder, N., Taxis, C., Reber, S., Maekawa, H., Moreno-Borchart, A., Doenges, G., Schwob, E., Schiebel, E., et al. (2004). A versatile toolbox for PCR-based tagging of yeast genes: new fluorescent proteins, more markers and promoter substitution cassettes. Yeast 21, 947–962. https://doi.org/10.1002/yea.1142.

Jessop, L., Rockmill, B., Roeder, G.S., and Lichten, M. (2006). Meiotic chromosome synapsis-promoting proteins antagonize the anti-crossover activity of sgs1. PLoS Genet. 2, e155. https://doi.org/10.1371/journal.pgen.0020155.

Johnson, F.B., Lombard, D.B., Neff, N.F., Mastrangelo, M.A., Dewolf, W., Ellis, N.A., Marciniak, R.A., Yin, Y., Jaenisch, R., and Guarente, L. (2000). Association of the Bloom syndrome protein with topoisomerase IIIalpha in somatic and meiotic cells. Cancer Res. 60, 1162–1167.

Jumper, J., Evans, R., Pritzel, A., Green, T., Figurnov, M., Ronneberger, O., Tunyasuvunakool, K., Bates, R., Žídek, A., Potapenko, A., et al. (2021). Highly accurate protein structure prediction with AlphaFold. Nature 596, 583–589. https://doi.org/10.1038/s41586-021-03819-2.

Kasaciunaite, K., Fettes, F., Levikova, M., Daldrop, P., Anand, R., Cejka, P., and Seidel, R. (2019). Competing interaction partners modulate the activity of Sgs1 helicase during DNA end resection. EMBO J. 38, e101516. https://doi.org/10.15252/embj.2019101516.

Knop, M., Siegers, K., Pereira, G., Zachariae, W., Winsor, B., Nasmyth, K., and Schiebel, E. (1999). Epitope tagging of yeast genes using a PCR-based strategy: more tags and improved practical routines. Yeast 15, 963–972. https://doi.org/10.1002/(SICI)1097-0061(199907)15:10B<963::AID-YEA399>3.0.CO;2-W.

Lagerwaard, I.M., Albanese, P., Jankevics, A., and Scheltema, R.A. (2022). Xlink Mapping and AnalySis (XMAS) -Smooth Integrative Modeling in ChimeraX.

Lam, I., and Keeney, S. (2014). Mechanism and regulation of meiotic recombination initiation. Cold Spring Harb. Perspect. Biol. 7, a016634. https://doi.org/10.1101/cshperspect.a016634.

LaRocque, J.R., Stark, J.M., Oh, J., Bojilova, E., Yusa, K., Horie, K., Takeda, J., and Jasin, M. (2011). Interhomolog recombination and loss of heterozygosity in wild-type and Bloom syndrome helicase (BLM)-deficient mammalian cells. Proc. Natl. Acad. Sci. U. S. A. 108, 11971–11976. https://doi.org/10.1073/pnas.1104421108.

Lynn, A., Soucek, R., and Börner, G.V. (2007). ZMM proteins during meiosis: crossover artists at work. Chromosome Res. 15, 591–605. https://doi.org/10.1007/s10577-007-1150-1.

Mazina, O.M., Mazin, A.V., Nakagawa, T., Kolodner, R.D., and Kowalczykowski, S.C. (2004). Saccharomyces cerevisiae Mer3 Helicase Stimulates 3t-5t Heteroduplex Extension by Rad51. Cell 117, 47–56. https://doi.org/10.1016/s0092-8674(04)00294-6.

Mercier, R., Jolivet, S., Vezon, D., Huppe, E., Chelysheva, L., Giovanni, M., Nogué, F., Doutriaux, M.-P., Horlow, C., Grelon, M., et al. (2005). Two meiotic crossover classes cohabit in Arabidopsis: one is dependent on MER3, whereas the other one is not. Curr. Biol. 15, 692–701. https://doi.org/10.1016/j.cub.2005.02.056.

Mullen, J.R., Nallaseth, F.S., Lan, Y.Q., Slagle, C.E., and Brill, S.J. (2005). Yeast Rmi1/Nce4 controls genome stability as a subunit of the Sgs1-Top3 complex. Mol. Cell. Biol. 25, 4476–4487. https://doi.org/10.1128/MCB.25.11.4476-4487.2005.

Mller, M.Q., Dreiocker, F., Ihling, C.H., Schäfer, M., and Sinz, (2010). Cleavable cross-linker for protein structure analysis: reliable identification of cross-linking products by tandem MS. Anal. Chem. 82, 6958–6968. https://doi.org/10.1021/ac101241t.

Nakagawa, T., and Kolodner, R.D. (2002). Saccharomyces cerevisiae Mer3 is a DNA helicase involved in meiotic crossing over. Mol. Cell. Biol. 22, 3281–3291. https://doi.org/10.1128/mcb.22.10.3281-3291.2002.

Nakagawa, T., Flores-Rozas, H., and Kolodner, R.D. (2001). The MER3 Helicase Involved in Meiotic Crossing Over Is Stimulated by Single-stranded DNA-binding Proteins and Unwinds DNA in the 3? to 5? Direction. J. Biol. Chem. 276, 31487–31493. https://doi.org/10.1074/jbc.M104003200.

Oh, S.D., Lao, J.P., Hwang, P.Y.-H., Taylor, A.F., Smith, G.R., and Hunter, N. (2007). BLM ortholog, Sgs1, prevents aberrant crossing-over by suppressing formation of multichromatid joint molecules. Cell 130, 259–272. https://doi.org/10.1016/j.cell.2007.05.035.

Pan, D., Brockmeyer, A., Mueller, F., Musacchio, A., and Bange, T. (2018). Simplified Protocol for Cross-linking Mass Spectrometry Using the MS-Cleavable Cross-linker DSBU with Efficient Cross-link Identification. Anal. Chem. 90, 10990–10999. https://doi.org/10.1021/acs.analchem.8b02593.

Pena, V., Jovin, S.M., Fabrizio, P., Orlowski, J., Bujnicki, J.M., Lührmann, R., and Wahl, M.C. (2009). Common design principles in the spliceosomal RNA helicase Brr2 and in the Hel308 DNA helicase. Mol. Cell 35, 454–466. https://doi.org/10.1016/j.molcel.2009.08.006.

Ponting, C.P. (2000). Proteins of the endoplasmic-reticulum-associated degradation pathway: domain detection and function prediction. Biochem. J 351 Pt 2, 527–535.

Pyatnitskaya, A., Borde, V., and Muyt, A.D. (2019). Crossing and zipping: molecular duties of the ZMM proteins in meiosis. Chromosoma 128, 181–198. https://doi.org/10.1007/s00412-019-00714-8.

Ranjha, L., Anand, R., and Cejka, P. (2014). The Saccharomyces cerevisiae Mlh1-Mlh3 heterodimer is an endonuclease that preferentially binds to Holliday junctions. J. Biol. Chem. 289, 5674–5686. https://doi.org/10.1074/jbc.M113.533810.

Schiffrin, B., Radford, S.E., Brockwell, D.J., and Calabrese, A.N. (2020). PyXlinkViewer: A flexible tool for visualization of protein chemical crosslinking data within the PyMOL molecular graphics system. Protein Sci. 29, 1851–1857. https://doi.org/10.1002/pro.3902.

Shinohara, M., Oh, S.D., Hunter, N., and Shinohara, A. (2008). Crossover assurance and crossover interference are distinctly regulated by the ZMM proteins during yeast meiosis. Nat. Genet. 40, 299–309. https://doi.org/10.1038/ng.83.

Storlazzi, A., Gargano, S., Ruprich-Robert, G., Falque, M., David, M., Kleckner, N., and Zickler, D. (2010). Recombination proteins mediate meiotic spatial chromosome organization and pairing. Cell 141, 94–106. https://doi.org/10.1016/j.cell.2010.02.041.

Tang, S., Wu, M.K.Y., Zhang, R., and Hunter, N. (2015). Pervasive and Essential Roles of the Top3-Rmi1 Decatenase Orchestrate Recombination and Facilitate Chromosome Segregation in Meiosis. Mol. Cell 57. https://doi.org/10.1016/j.molcel.2015.01.021.

Vernekar, D.V., Reginato, G., Adam, C., Ranjha, L., Dingli, F., Marsolier, M.-C., Loew, D., Guis, R., Llorente, B., Cejka, P., et al. (2021). The Pif1 helicase is actively inhibited during meiotic recombination which restrains gene conversion tract length. Nucleic Acids Res. 49, 4522–4533. https://doi.org/10.1093/nar/gkab232.

Wang, T.-F., and Kung, W.-M. (2002). Supercomplex formation between Mlh1-Mlh3 and Sgs1-Top3 heterocomplexes in meiotic yeast cells. Biochem. Biophys. Res. Commun. 296, 949–953. https://doi.org/10.1016/s0006-291x(02)02034-x.

Wang, J., Zhang, W., Jiang, H., Wu, B.-L., and Primary Ovarian Insufficiency Collaboration (2014). Mutations in HFM1 in recessive primary ovarian insufficiency. N. Engl. J. Med. 370, 972–974. https://doi.org/10.1056/NEJMc1310150.

Wang, K., Tang, D., Wang, M., Lu, J., Yu, H., Liu, J., Qian, B., Gong, Z., Wang, X., Chen, J., et al. (2009). MER3 is required for normal meiotic crossover formation, but not for presynaptic alignment in rice. J. Cell Sci. 122, 2055–2063. https://doi.org/10.1242/jcs.049080.

Weissmann, F., Petzold, G., VanderLinden, R., Veld, P.J.H.I. ‘t, Brown, N.G., Lampert, F., Westermann, S., Stark, H., Schulman, B.A., and Peters, J.-M. (2016). biGBac enables rapid gene assembly for the expression of large multisubunit protein complexes. Proc. Natl. Acad. Sci. U. S. A. 113, E2564 9. https://doi.org/10.1073/pnas.1604935113.

Wild, P., Susperregui, A., Piazza, I., Dörig, C., Oke, A., Arter, M., Yamaguchi, M., Hilditch, A.T., Vuina, K., Chan, K.C., et al. (2019). Network Rewiring of Homologous Recombination Enzymes during Mitotic Proliferation and Meiosis. Mol. Cell 75, 859-874.e4. https://doi.org/10.1016/j.molcel.2019.06.022.

Yadav, V.K., and Claeys Bouuaert, C. (2021). Mechanism and Control of Meiotic DNA Double-Strand Break Formation in S. cerevisiae. Front Cell Dev Biol 9, 642737. https://doi.org/10.3389/fcell.2021.642737.

